# Growth-regulated Hsp70 phosphorylation regulates stress responses and prion maintenance

**DOI:** 10.1101/759241

**Authors:** Chung-Hsuan Kao, Seung Ryu, Min J. Kim, Xuemei Wen, Oshadi Wimalarathne, Tanya T. Paull

## Abstract

Maintenance of protein homeostasis in eukaryotes during normal growth and stress conditions requires the functions of Hsp70 chaperones and associated co-chaperones. Here we investigate an evolutionarily-conserved serine phosphorylation that occurs at the site of communication between the nucleotide-binding and substrate-binding domains of Hsp70. Ser151 phosphorylation in yeast Hsp70 (Ssa1) is promoted by cyclin-dependent kinase (Cdk1) during normal growth and dramatically affects heat shock responses, a function conserved with Hsc70 S153 phosphorylation in human cells. Phospho-mimic forms of Ssa1 (S151D) also fail to relocalize in response to starvation conditions, do not associate *in vivo* with Hsp40 co-chaperones, Ydj1 and Sis1, and do not catalyze refolding of denatured proteins *in vitro* in cooperation with Ydj1 and Hsp104. S151 phosphorylation strongly promotes survival of heavy metal exposure and reduces Sup35-dependent *[PSI^+^]* prion activity, however, consistent with proposed roles for Ssa1 and Hsp104 in generating self-nucleating seeds of misfolded proteins. Taken together, these results suggest that Cdk1 downregulates Hsp70 function during periods of active growth, reducing propagation of aggregated proteins despite potential costs to overall chaperone efficiency.

---

Protein homeostasis encompasses a network of processes which maintain the functionality of proteins in the cellular environment (Kim *et al*, 2013). Nascent polypeptides fold into stable, tertiary structures during and after translation; however, mammalian protein biosynthesis often leads to incorrectly folded proteins. It is estimated that 30 – 50% of mammalian proteins are in misfolded or metastable states with partially ordered structures (Balchin *et al*, 2016; Saibil, 2013). These forms of proteins may lead to the accumulation of aggregates, including amorphous, oligomeric, and fibrillar structures, which are toxic to cells (Balchin *et al*, 2016). In order to maintain efficient protein folding and prevent protein aggregation, cells have evolved a network of chaperone proteins as a surveillance system for proteome integrity. Disruption of these networks can cause a number of human disorders, including Alzheimer’s, Parkinson’s, and prion-related diseases (Soto, 2003). While the activities of many members of the chaperone family are well studied, the regulation of these critical enzymes during specific stress conditions is not completely understood.

Chaperones have been classified into Hsp70, Hsp90, Hsp100, Hsp110, Hsp40, and small HSP families (Genest *et al*, 2019). The canonical function of Hsp70s and Hsp90 chaperones are to recognize unfolded protein clients, promote correct folding, and release them after the protein folding is completed. Protein folding requires an ATP/ADP exchange cycle and the assistance of co-chaperones (Saibil, 2013). Two major co-chaperones, Hsp110 (nucleotide exchange factors, NEFs) and Hsp40 (DNAJ-related proteins), cooperate with Hsp70s and regulate the exchange between ATP-bound and ADP-bound states (Kampinga & Craig, 2010). Hsp70 proteins also work as a hub connecting with other chaperones to facilitate translocation between cellular compartments, regulation of newly synthesized proteins, and sequestration and degradation of protein aggregates (Chen *et al*, 2011).

Hsp70 proteins are highly conserved in all species (Daugaard *et al*, 2007). In *Saccharomyces cerevisiae*, there are four functionally and structurally redundant Hsp70 proteins: Ssa1-4. While Ssa1 and Ssa2 are constitutively expressed similar to human Hsc70, Ssa3 and Ssa4 are induced by heat shock and other forms of stress similar to human Hsp70 (Verghese *et al*, 2012). Removal of Ssa1-4 is lethal in yeast, but constitutive expression of any single gene can rescue this lethality (Werner-Washburne *et al*, 1987; Hasin *et al*, 2014). Hsp70 orthologs in eukaryotes are targets of many post-translational modifications, including numerous phosphorylation events (Cloutier & Coulombe, 2013). Yeast Ssa1 has two phosphorylation hotspots: one in the N-terminal nucleotide binding domain (NBD) and C-terminal substrate binding domain (SBD). Some of these phosphorylation events have been characterized and shown to regulate HSP70/SSA-dependent functions including heat shock responses, polysome association, protein refolding, and protein disaggregation (Beltrao *et al*, 2012). There have also been numerous other post-translational modifications observed in HSP70 proteins in eukaryotes (Cloutier & Coulombe, 2013), but most of these are uncharacterized.

Here we investigated a conserved phosphorylation site in the ATPase domain of HSP70 enzymes and found that growth-dependent modification of this site controls HSP70 function in budding yeast and in mammalian cells. Phosphorylation of S151 (S151) on yeast Ssa1 regulates its association with co-chaperone and chaperone partners and modulates ATP hydrolysis and protein disaggregation. During heat stress or nutrition deficiency, phosphorylation of S151 plays an important role in regulating the Hsp104 disaggregation machinery and stress granule formation, respectively. In addition, we find that the phosphorylation status of S151 dictates the efficiency of Sup35 prion maintenance as well as survival of heavy metal exposure. Based on this evidence, we propose that Ssa1 S151 phosphorylation is critical for the dynamic association of the protein with co-chaperones and clients during stress responses and may play a role in the regulation of misfolded proteins in coordination with nutritional states.

## Results

### Ssa1 S151 phosphorylation affects cell growth and thermal stability

Multiple phosphorylated serine/threonine residues have been identified on HSP70 proteins (Cloutier & Coulombe, 2013; Albuquerque *et al*, 2008; Holt *et al*, 2009); however, the potential functions and regulatory roles of these modifications on Hsp70 are not fully understood. To identify and characterize these sites, we analyzed Ssa1 phosphorylation in budding yeast and Hsc70 phosphorylation in human cells by quantitative mass spectrometry. In our analysis of post-translational modifications, we detected and confirmed Ssa1 and Hsc70 phosphorylation at S151 and S153 in yeast and humans, respectively, among many other modifications (Table S1 and S2). The S151 residue is highly conserved among several Hsp70 family members in *S. cerevisiae* and higher eukaryotes (Fig. 1A). The four abundant cytosolic HSP70 proteins (Ssa1-4) and yeast mitochondrial HSP70 proteins (Ssc1 and Ssq1) all have a serine at a.a.151 in the nucleotide-binding domain (NBD) whereas most bacterial and archaeabacterial HSP70 proteins have an alanine at this position.

**Figure 1.**
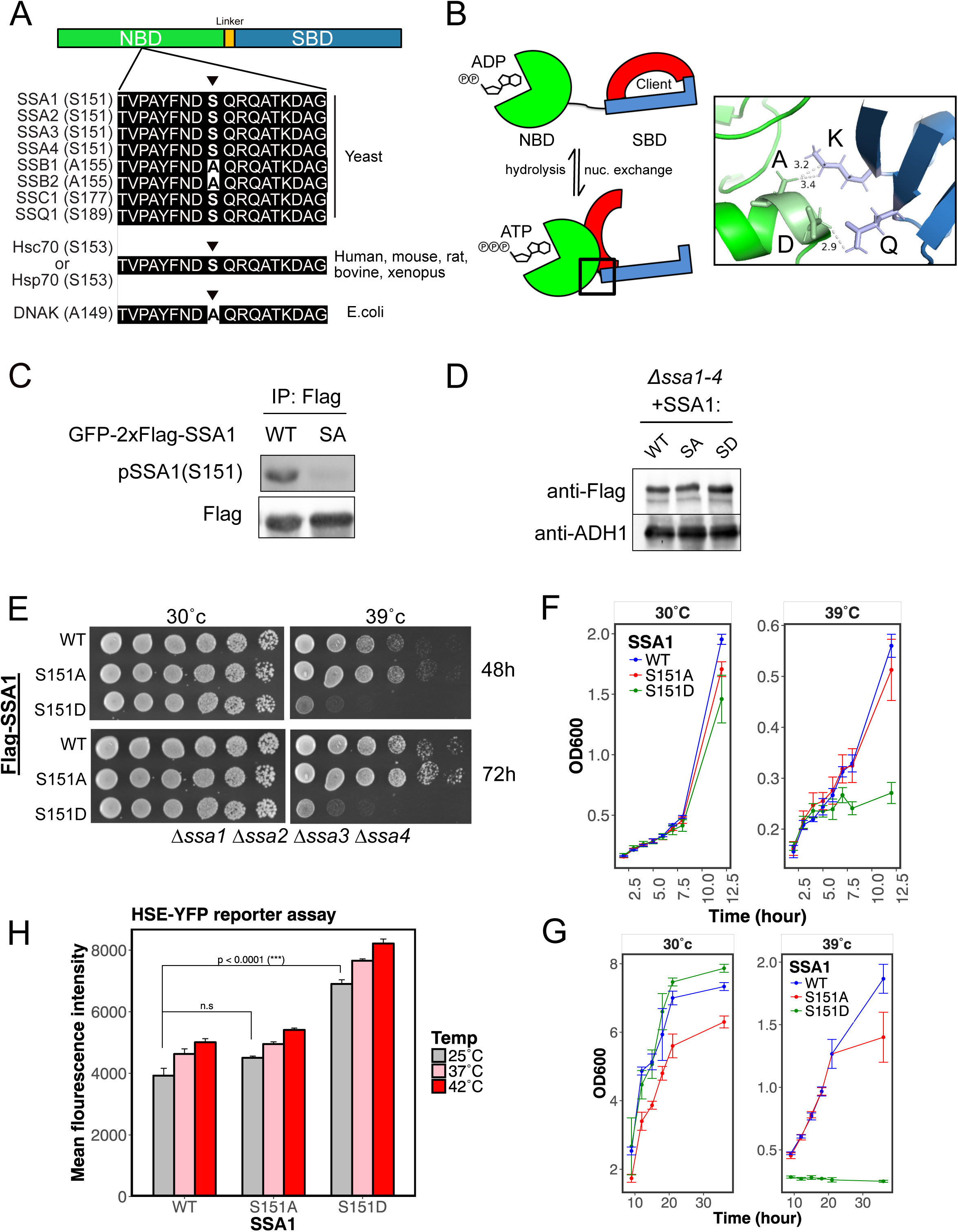
Ssa1 S151 occurs in budding yeast and affects survival of heat shock. (A) Alignment of Hsp70 protein sequences in the region surrounding S151 in the NBD in prokaryotic and eukaryotic cells as indicated. (B) The S151 residue of Ssa1 in the NBD is located close to the interaction site with the SBD in the ATP-bound state. Left panel: ATP hydrolysis and nucleotide exchange is postulated to regulate structural conformation changes in Hsp70 proteins. In the ADP-bound state, the NBD (green) is in an open configuration, connected to the SBD (red: alpha-helical lid; blue: beta-sheet pocket) via a flexible linker. In the ATP-bound state, NBD and SBD undergo a conformational change to interact in a closed configuration. Right panel: In DnaK, A149 and D148 of the NBD are in close contact with K452 and E442 of the SBD (PDB: 4B9Q) (Kityk *et al*, 2012). (C) GFP-Flag-Ssa1 (WT) or GFP-Flag-Ssa1 S151A (S151A) were expressed in a *ssa1 ssa2 ssa3 ssa4* deletion (*Δssa1-4*) strain. GFP-Flag-Ssa1 was isolated by immunoprecipitation and analyzed by quantitative Western blotting with anti-phospho-Ssa1 S151 and anti-Flag antibodies. (D) Total lysates from *Δssa1-4* yeast cells expressing Flag-Ssa1 (WT), Flag-Ssa1 S151A (S151A), or Flag-Ssa1 S151D (S151D) were analyzed by wester blotting for Flag or ADH1 as a loading control. (E) *Δssa1-4* yeast cells expressing Flag-Ssa1 (WT), Flag-Ssa1 S151A (S151A), or Flag-Ssa1 S151D (S151D) were spotted in fivefold serial dilutions and exposed to 30 or 39°C for 48 or 72 hr. (F and G) Growth of Δ*ssa1-4* yeast cells expressing Flag-Ssa1 (WT), Flag-Ssa1 S151A (S151A), or Flag-Ssa1 S151D (S151D) was monitored at 30°C or 39°C as indicated. Growth curve is measured in log phase (F) and stationary (G) phase by OD600. 3 biological replicates were performed and error bars represent standard deviation. (H) Δ*ssa1-4* yeast cells with an integrated HSE-YFP reporter and expressing Flag-Ssa1 (WT), Flag-Ssa1 S151A (S151A), or Flag-Ssa1 S151D (S151D) were exposed to varying temperatures as indicated for 30 min. YFP signal was measured by flow cytometer. 3 biological replicates were performed with at least 10,000 cells per measurement; error bars represent standard deviation.

In Hsc70/Hsp70 proteins, the substrate-binding domain (SBD) docks with the NBD in the ATP-bound state where the complex has low affinity for its clients (Kityk *et al*, 2012; Saibil, 2013) (example of the ATP-bound form of DnaK (*E. coli* homologue of mammalian Hsp70) shown in Fig. 1B (Kityk *et al*, 2012). With ATP hydrolysis, the SBD releases from this conformation while connected to the NBD via a flexible linker in the ADP-bound state, consistent with predictions of dynamic motion of Hsp70 between different states (Flaherty *et al*, 1990; Zhu *et al*, 1996). Interestingly, the Alanine 149 (A149) residue of DnaK (corresponding to the S151 residue of yeast Ssa1 or S153 residue of human Hsc70/Hsp70) is located close to the interface between the NBD and SBD (Fig. 1B). Based on structures of DnaK and yeast Sse1 (Kityk *et al*, 2012; Liu & Hendrickson, 2007), S151/153 would only be ∼3Å away from Lysine 452 (K452) of the SBD in the ATP-bound state, indicating a possible role for this modification in regulating associations between the NBD and the SBD.

Interestingly, S153 in vertebrate Hsc70 orthologs is part of an (S/T)Q motif, matching the consensus for PI-3 Kinase-like Kinases (PIKK) that are targets of the ATM and ATR protein kinases in the DNA damage response (Kim *et al*, 1999; O’Neill *et al*, 2000). Hsc70 S153 was also found to be phosphorylated in a global study of DNA damage-induced phosphorylation (Matsuoka *et al*, 2007). In yeast, S151 in SSA1 also resides in an (S/T)Q PIKK motif; however, the role of SSA1 S151 phosphorylation in DNA damage responses is not clear. Our observation of SSA1 S151 phosphorylation as well as previous documentation of this modification by global phosphorylation studies (Beltrao *et al*, 2012; Albuquerque *et al*, 2008) suggested that one of the PIKK kinases might use Hsp70 modification to regulate downstream DNA damage repair pathways or canonical Hsp70 functions. To investigate this possibility, we generated a custom antibody directed against phospho-S151 and monitored phosphorylation of recombinant GFP-Flag-Ssa1 expressed in budding yeast and isolated by immunoprecipitation followed by western blot of the tagged protein from normally growing cells. The result shows that the antibody recognizes wild-type Ssa1 but not Ssa1 S151A (Fig. 1C), indicating that the antibody is specific for the S151 residue and that the modification occurs during normal growth conditions. We did not observe any increase in phosphorylation with DNA damaging agents, however (data not shown), which would be expected if this modification was involved in DNA damage responses.

In order to investigate the role of HSP70 phosphorylation at S151, we expressed Flag-tagged wild-type Ssa1, a non-phosphorylatable mutant Ssa1 (S151A) or a phospho-mimetic mutant Ssa1 (S151D) under the control of the Ssa1 natural promoter on a 2 micron plasmid in a yeast strain lacking all four Ssa proteins (Fig. 1D)(Jaiswal *et al*, 2011). Growth of wild-type and mutant strains were evaluated using a serial spot dilution assay on solid media (Fig. 1E) or a growth curve in liquid culture (Fig. 1F, G). Both assays indicate that the wild-type, S151A, and S151D cells grow similarly at 30°C but the S151D cells are hypersensitive to high temperature (39°C). Therefore, the phospho-mimetic S151D mutant can be considered a temperature-sensitive mutant. Interestingly, we observed that the cells expressing the S151A mutant exhibit a delay in growth as cells approach stationary phase, whereas the cells expressing S151D grow to higher density than wild-type during this period (Fig. 1F).

It is possible that S151D cells exhibit sensitivity to heat shock because of alterations in heat shock factor 1 (Hsf1) regulation (Trotter *et al*, 2001). In wild-type *S. cerevisiae* cells, the transcription factor Hsf1 is repressed by Ssa1/2, which binds to Hsf1, preventing its association with heat shock elements (HSEs) (Sorger & Pelham, 1988). Heat shock generates misfolded proteins that compete with Hsf1 for the binding of Ssa1/2, which releases Hsf1 to transcribe HSE-driven genes that generate higher levels of chaperones and repress ribosomal proteins (Trotter *et al*, 2001; Rowley *et al*, 1993; Zheng *et al*, 2016). To test whether Ssa1 S151 phosphorylation regulates HSF1 activation, we integrated a yellow fluorescence protein (YFP) reporter regulated by HSEs (Zheng *et al*, 2016) into the genome of our strains and used flow cytometry analysis to monitor HSF1 activity. In the wild-type and S151A strains, exposure to heat (37°C and 42°C) increased the yield of YFP relative to 25°C (Fig. 1H) as previously reported (Zheng *et al*, 2016). Cells expressing S151D Ssa1 exhibited higher HSF1 activation at high temperatures but were also elevated at 25°C compared to wild-type and S151A cells. Consistent with this finding of basal de-repression of Hsf1, cells expressing S151D Ssa1 under normal growth conditions show higher levels of expression of several other heat shock proteins (Fig. S1) and also slight repression of ribosomal proteins (Fig. S2).

### Ssa1 S151D cells accumulate more heat-induced protein aggregates

The impaired heat shock survival of yeast cells expressing Ssa1 S151D protein (Fig. 1D-F) might be related to the accumulation of heat-induced protein aggregates, as Hsp70 proteins have been shown to be critical for dispersal of these toxic products (Mogk *et al*, 2018). To test this, we collected protein aggregates by separating detergent-insoluble proteins from detergent-soluble proteins using a previously described method (Koplin *et al*, 2010). We then compared overall levels of detergent-insoluble proteins from wild-type, S151A, and S151D cells at 30°C or 42°C by coomassie staining of the aggregates on SDS-PAGE gels (Fig. 2A). Interestingly, cells expressing Ssa1 S151D tended to form a higher basal level of total endogenous aggregates than wild-type and Ssa1 S151A cells at 30°C (“C”). Similarly, Ssa1 S151D cells produced more heat-induced total aggregates compared to wild-type and S151A cells at 42°C (“Hs”, Fig. 2A). The results are consistent with the growth data above in that S151D cells accumulate more detergent-insoluble proteins than wild-type and S151A cells during heat exposure. In contrast, the S151A-expressing cells showed lower levels of protein aggregates compared to wild-type Ssa1-expressing cells, with or without heat treatment, suggesting a higher efficiency of aggregate removal compared to wild-type cells. Arsenite treatment, used here as a general form of oxidative stress, yielded similar levels of aggregates in all strains.

**Figure 2.**
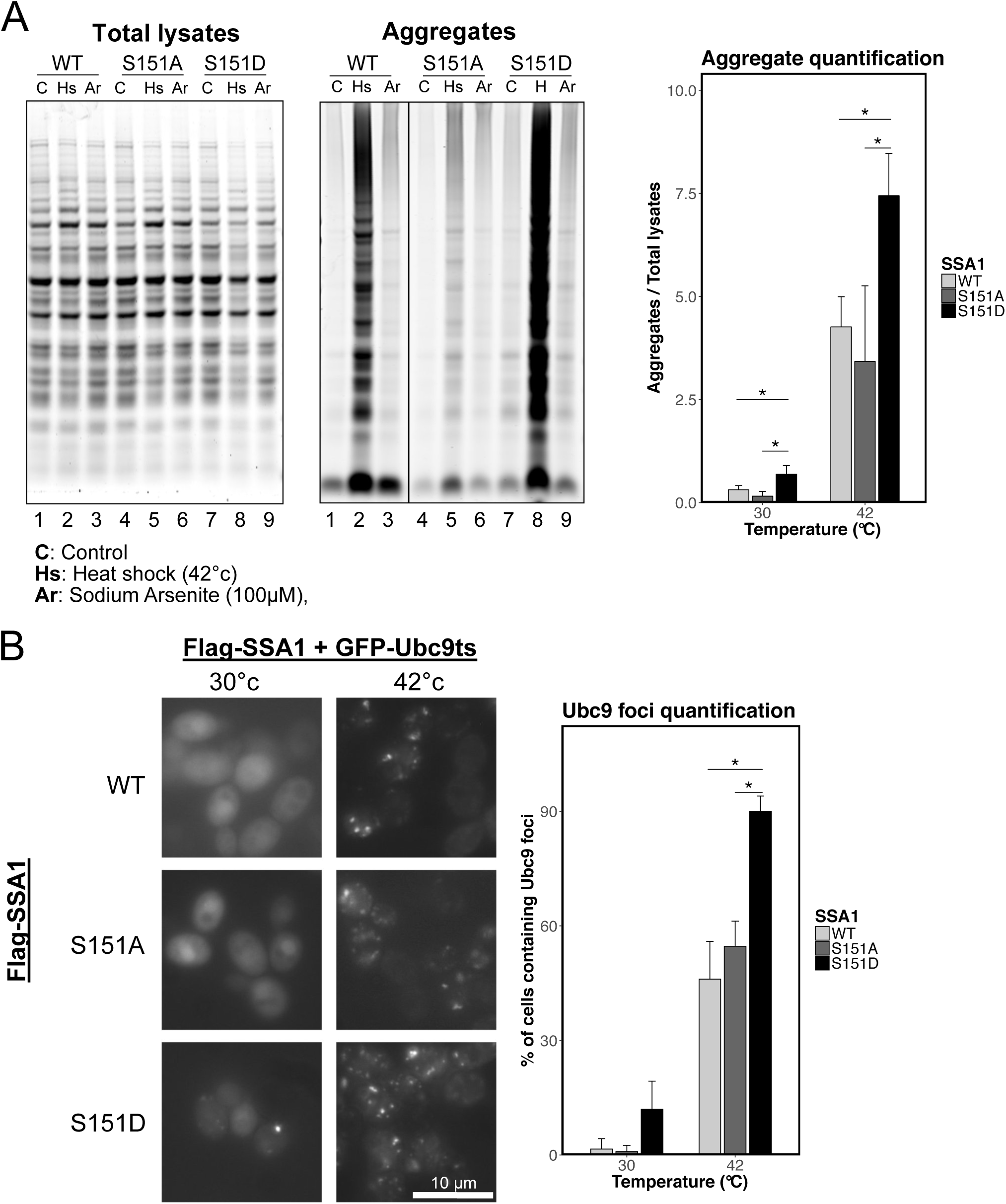
Ssa1 S151 phosphorylation promotes heat-induced protein aggregation. (A) Left and middle panels: *Δssa1-4* yeast cells expressing Flag-Ssa1 (WT), Flag-Ssa1 S151A (S151A), or Flag-Ssa1 S151D (S151D) were treated with heat shock (42°C, “Hs”), or Sodium Arsenite (100 μM, “Ar”) for 60 min. Protein aggregates were isolated (see Materials and Methods) and separated by SDS-PAGE with total lysates. Right panel: Quantification of the levels of aggregated proteins divided by the total protein levels. 3 biological replicates were performed and error bars represent standard deviation. * indicates p < 0.05 by student’s 2-tailed T test. (B) Left panel: Δ*ssa1-4* yeast cells expressing GFP-Ubc9ts as well as Flag-Ssa1 (WT), Flag-Ssa1 S151A (S151A), or Flag-Ssa1 S151D (S151D) were treated with 30°C or 42°C for 30 min and analyzed by immunofluorescence microscope for Ubc9 foci. Right panel: Quantification of Ubc9 foci was performed by counting the number of GFP-positive cells containing at least one GFP focus per cell divided by the total number of GFP-positive cells. 3 biological replicates were performed and error bars represent standard deviation. * indicates p < 0.05 by student 2-tailed T test.

We also used the well-established aggregate reporter Ubc9 Y68L, a temperature-sensitive allele of the SUMO-conjugating enzyme Ubc9 (Ubc9^ts^), to measure protein aggregates in vivo (Kaganovich *et al*, 2008; Betting & Seufert, 1996). Our results showed that cells expressing Ssa1 S151D form more endogenous GFP-Ubc9 aggregates (GFP foci) at 30°C and also show nearly 2-fold higher levels of heat-induced GFP foci compared to the wild-type and S151A cells (Fig. 2B). Taken together, these data suggest that Ssa1 S151 phosphorylation affects the cellular responses to heat and protein homeostasis stress.

### Ssa1 S151 is phosphorylated by cyclin-dependent kinase Cdk1

Although we initiated our study of S151 modification with the idea that PIKK enzymes might regulate HSP70 function, analysis of S151 phosphorylation in strains deficient in the yeast PIKK enzymes (*mec1 tel1* strains) did not support this hypothesis (data not shown). A recent study of other Ssa1 modifications showed that T36 phosphorylation on Ssa1 by yeast Cdk1 and Pho85 is important for G_1_/S cell cycle control and suggested that Ssa1 is physically associated with Cdk1 and Pho85 (Truman *et al*, 2012). Considering this precedent, we hypothesized that cyclin-dependent kinase might phosphorylate Ssa1 S151. To determine if Cdk1 catalyzes Ssa1 S151 phosphorylation, we expressed galactose-inducible Cdk1(Cdc28) in GFP-Flag-SSA1-expressing cells and analyzed the phosphorylation status of S151 in Ssa1 immunoprecipitated with anti-Flag antibody. The result shows that Cdk1 overexpression increases Ssa1 S151 phosphorylation signal 3-fold, with Ssa1 S151A as a negative control (Fig. 3A).

**Figure 3.**
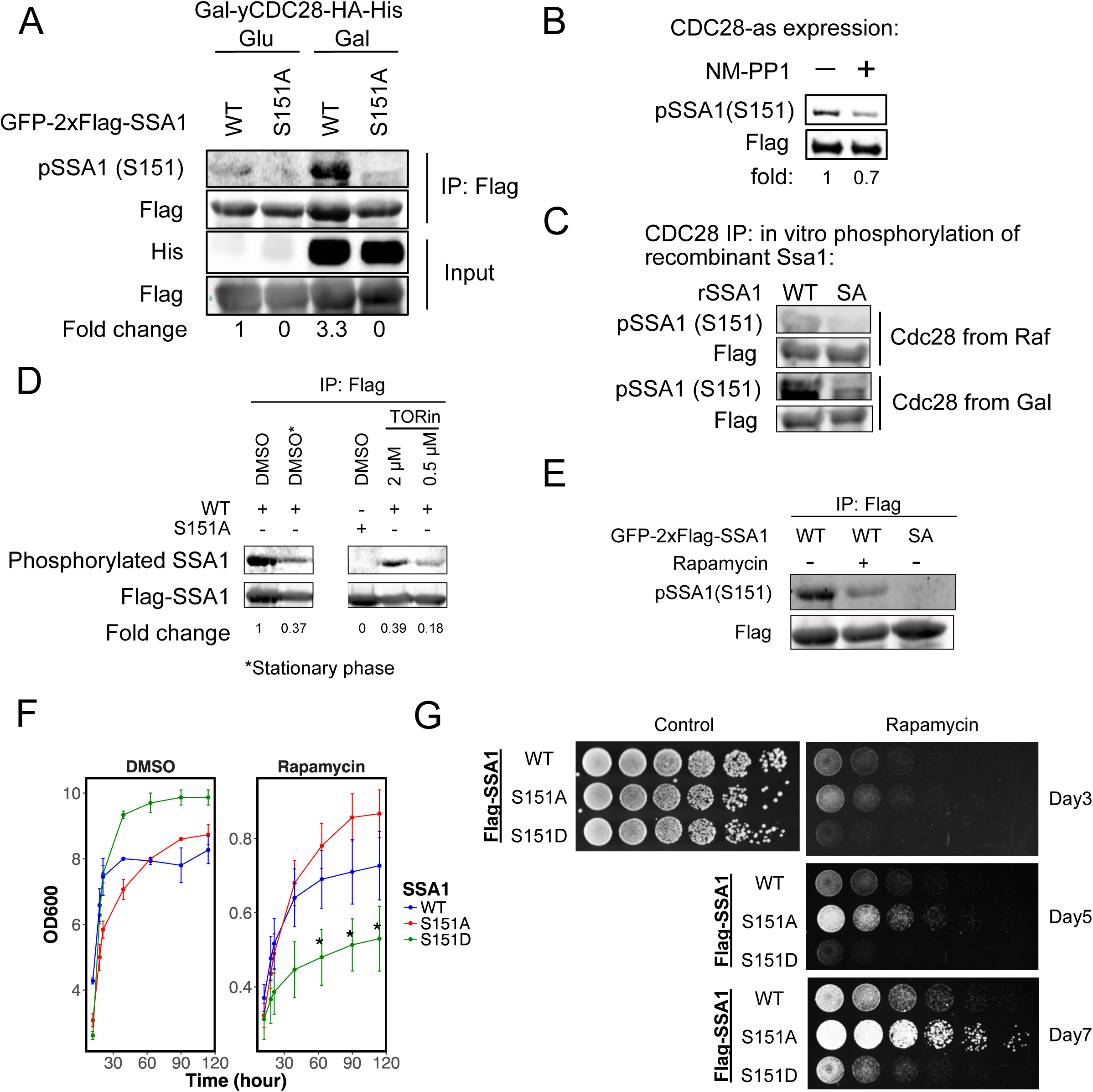
Ssa1 S151 phosphorylation is mediated by Cdk1 and regulated by the TOR pathway. (A) Δ*ssa1-4* yeast cells expressing GFP-Flag-Ssa1 (WT), or GFP-Flag-Ssa1 S151A (S151A) as well as galactose-inducible HA-His-tagged Cdk1 (Gal-yCDC28-HA-His) were grown in glucose (Glu) or galactose (Gal). Ssa1 was isolated by immunoprecipitation and analyzed by Western blotting with anti-phospho-Ssa1 S151, anti-His and anti-Flag antibodies. (B) Cdk1-as1 cells expressing GFP-Flag-Ssa1 (WT), or GFP-Flag-Ssa1 S151A (S151A) were treated with NM-PP1 (10 uM) for 3 hr. Ssa1 was isolated by immunoprecipitation and analyzed by Western blotting with anti-phospho-Ssa1 S151 and anti-Flag antibodies. (C) Recombinant Flag-tagged Ssa1 (WT) or Ssa1 S151A (S151A) was incubated with HA-His-tagged Cdk1 isolated from *ssa1-4*Δ yeast cells expressing galactose inducible HA-His-tagged Cdk1 (Gal-yCDC28-HA-His) treated with raffinose (Raf) or galactose (Gal). Phosphorylation was analyzed by Western blotting with anti-phospho-Ssa1 S151 and Flag antibodies. Arrow indicates phosphorylated species. (D) Δ*ssa1-4* yeast cells expressing GFP-Flag-Ssa1 (WT), or GFP-Flag-Ssa1 S151A (S151A) were treated with TORin (2 μM or 0.5 μM) for 30 min. Ssa1 was isolated by immunoprecipitation and analyzed by Western blotting with anti-phospho-Ssa1 S151 and anti-Flag antibodies. (E) Δ*ssa1-4* yeast cells expressing GFP-Flag-Ssa1 (WT), or GFP-Flag-Ssa1 S151A (S151A) were treated with rapamycin (200 ng/mL) for 30 min. Ssa1 was isolated by immunoprecipitation and analyzed by quantitative Western blotting with anti-phospho-Ssa1 S151 and anti-Flag antibodies. (F) Δ*ssa1-4* yeast cells expressing Flag-Ssa1 (WT), Flag-Ssa1 S151A (S151A), or Flag-Ssa1 S151D (S151D) were monitored for growth over time in the absence or presence of rapamycin (100 ng/mL). (G) Δ*ssa1-4* yeast cells expressing Flag-Ssa1 (WT), Flag-Ssa1 S151A (S151A), or Flag-Ssa1 S151D (S151D) were spotted in fivefold serial dilution on control or rapamycin plates (100 ng/mL) as indicated.

We also utilized yeast cells expressing Cdk1-as1, an analog-sensitive version of Cdk1 (Bishop *et al*, 2000), to evaluate Ssa1 S151 phosphorylation. In this strain, Cdk1 activity can be downregulated by the addition of 1-NM-PP1 analogs that uniquely block the analog-sensitive Cdk1. We overexpressed GFP-Flag-SSA1 in the Cdk1-as1 strain and immunoprecipitated Ssa1 after 1-NM-PP1 treatment, finding that Ssa1 phosphorylation at S151 was partially decreased under these conditions (Fig. 3B). Lastly, to confirm the phosphorylation in vitro, we purified galacotose-inducible His-HA-tagged Cdk1 enzyme and evaluated its ability to phosphorylate recombinant wild-type Ssa1 purified from *E. coli*. In comparison to the purified Cdk1 from raffinose cultures, immunoprecipitated material from galactose-induced cultures produced 1.5 to 2--fold higher levels of Ssa1 S151 phosphorylation (Fig. 3C). In addition, the S151 site was found in a global survey of Cdk1 targets in yeast, although the change in phosphorylation induced by cell cycle synchronization in this study was modest (Holt *et al*, 2009). Taken together, these data suggest that Cdk1 phosphorylates Ssa1 at S151 in vivo, although we cannot exclude the possibility of other kinases modifying the site in addition to Cdk1.

### Starvation conditions reduce S151 phosphorylation

The TOR pathway participates in protein homeostasis through the connection between TOR-mediated nutrient signaling and chaperone-mediated stress responses (Conn & Qian, 2011). Here we tested the effects of stationary phase, TORin (a TOR1 and TOR2 inhibitor)(Huang *et al*, 2017), as well as rapamycin and found that all of these treatments substantially reduce Ssa1 phosphorylation at S151 (Fig. 3D, E). These results are consistent with our observation of Cdk1 control of this phosphorylation since TOR inhibition blocks cell cycle progression and Cdk1 activity is low in stationary phase (Barbet *et al*, 1996; Hartwell *et al*, 1974; Mendenhall *et al*, 1987). We also tested the growth of wild-type and mutant strains under TOR inhibition conditions using a growth curve in liquid culture (Fig. 3F) or a serial spot dilution assay on solid media (Fig. 3G). Both assays indicated that S151D cells exhibit delayed growth in a TOR-inhibited environment, particularly in high-density cultures as cells approach stationary phase. Interestingly, S151A cells grew to higher densities than wild-type and S151D cells during long-term rapamycin treatment (Fig. 3F, G). Thus, reduction of S151 phosphorylation is necessary for growth and survival under conditions of TOR inhibition.

Translation inhibition is a common response to many types of environmental stress (Cherkasov *et al*, 2013). Cellular RNA-containing granules form during nutrient deprivation and during stationary phase and serve to protect mRNAs as well as to promote translation re-initiation and elongation after stress conditions are resolved. In budding yeast, Ssa proteins and Hsp40 co-chaperones are important for the formation of cytoplasmic RNA-protein (RNP) granules (stress granule and P-bodies)(Walters *et al*, 2015). Here we investigated the efficiency of RNP granule formation by examining GFP-SSA1 foci formation during glucose deprivation and stationary phase. The results showed that cells expressing Ssa1 S151D completely failed to form RNP granules during glucose deprivation (Fig. 4A). We also examined the behavior of GFP-Ssa1 in stationary phase during which the chaperone also is known to form discrete foci (Narayanaswamy *et al*, 2009). We observed a lower density of RNP granules in both S151A and S151D cells compared to wild-type cells, although S151D expression reduced foci to a greater extent than S151A (Fig. 4B). Thus, reorganization of Ssa1 into stress granules is sensitive to the phosphorylation status of S151.

**Figure 4.**
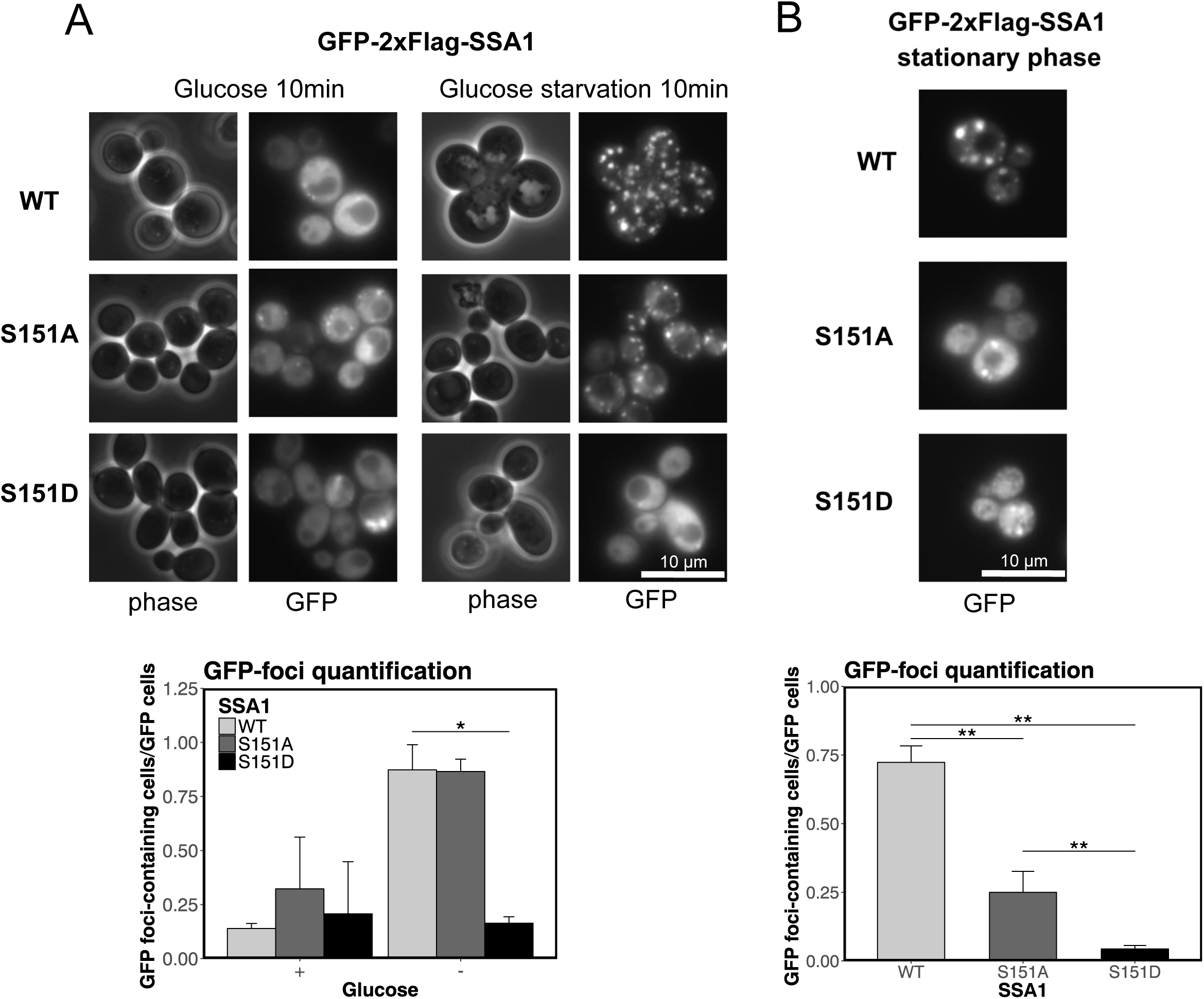
Ssa1 S151 phosphorylation regulates chaperone localization in response to stress. (A) Top panel: Δ*ssa1-4* yeast cells expressing GFP-Flag-Ssa1 (WT), GFP-Flag-Ssa1 S151A (S151A), or GFP-Flag-Ssa1 S151D (S151D) were incubated with or without 2% glucose for 10 min and analyzed by fluorescence microscopy. Bottom panel: Quantification of GFP foci was performed by counting the number of cells containing at least 1 GFP focus divided by the total number of GFP-positive cells. 3 biological replicates were performed with at least 50 cells counted per measurement; error bars represent standard deviation. * indicates p < 0.05 by student’s 2-tailed T test. (B) Top panel: Δ*ssa1-4* yeast cells expressing GFP-Flag-Ssa1 (WT), GFP-Flag-Ssa1 S151A (S151A), or GFP-Flag-Ssa1 S151D (S151D) were incubated at 30°C for 3 days to reach saturation and analyzed by fluorescence microscopy for GFP foci. Bottom panel: Quantification of GFP foci was performed by counting the number of cells containing at least 1 GFP focus divided by the total number of GFP-positive cells. 3 biological replicates were performed with at least 50 cells counted per measurement; error bars represent standard deviation. ** indicates p < 0.05 by student 2-tailed T test.

### S151 phosphorylation regulates the interactome of Ssa1

Hsp70 orthologs participate in a wide range of cellular processes through their ATP-dependent cycles of client recognition and protein folding (Clerico *et al*, 2015). In order to investigate the impact of Ssa1 S151 phosphorylation on the global interactome of Ssa1, we purified GFP-Flag-tagged Ssa1 wild-type, Ssa1 (S151A), and Ssa1 (S151D) proteins from yeast cells during exponential growth and compared their binding partners by quantitative liquid chromatography-tandem mass spectrometry (LC-MS/MS). We detected 2006 proteins in lysates and immunoprecipitates and used untagged Ssa1 for immunoprecipitation as a negative control. The bound proteins included known co-chaperones and chaperone-associated factors as well as other proteins that may be clients. We identified 46 proteins significantly altered in their binding to S151A compared to wild-type chaperone, while 57 proteins are altered in cells expressing S151D, comparing 3 biological replicates from each strain (Fig. 5A-C, Table S3). These results suggested that the Ssa1 S151 phosphorylation broadly influences the binding between Ssa1 and cellular factors.

**Figure 5.**
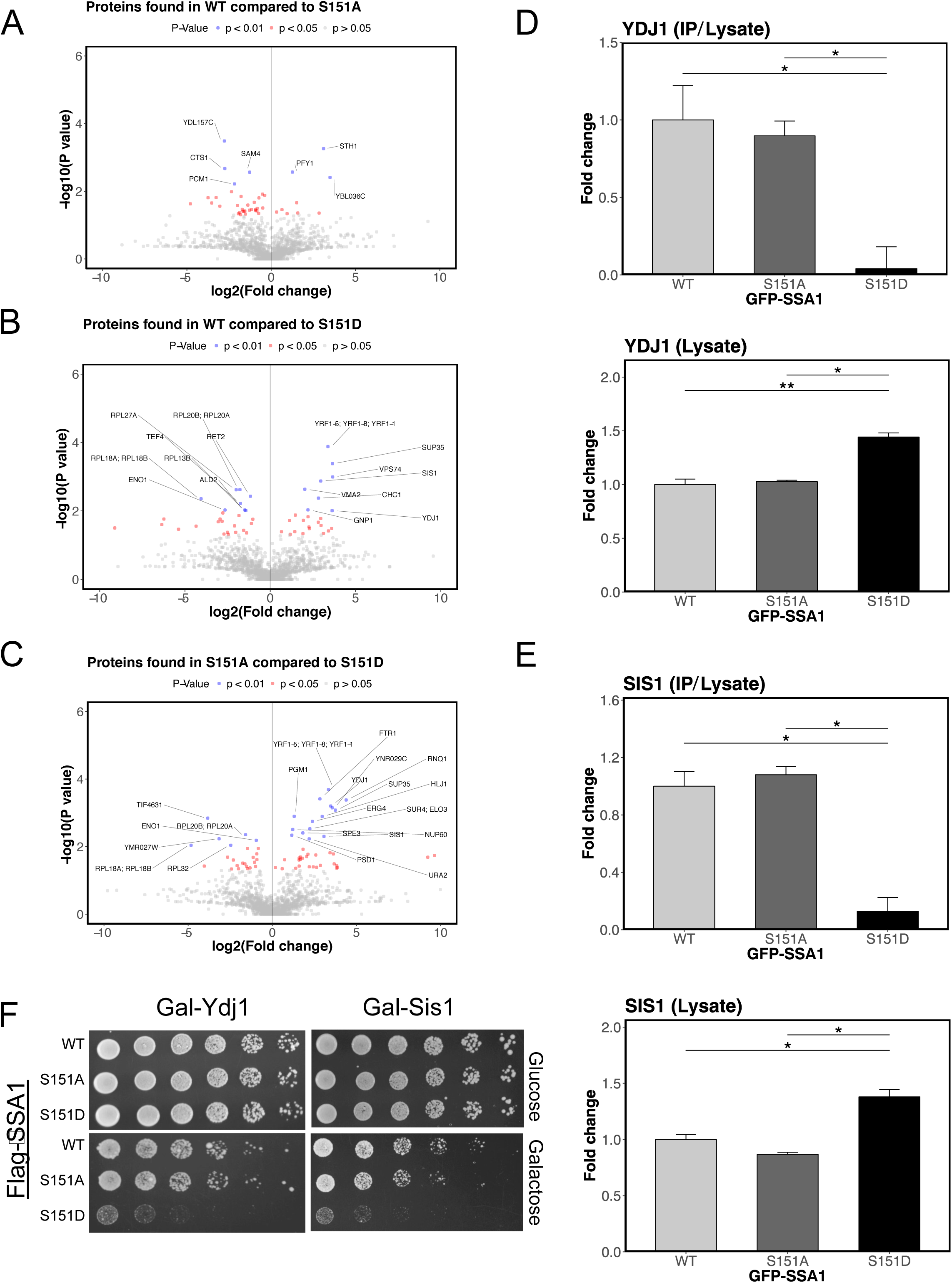
S151 phosphorylation regulates the interactome of Ssa1. Δ*ssa1-4* yeast cells expressing GFP-Flag-Ssa1 (WT), GFP-Flag-Ssa1 S151A (S151A), or GFP-Flag-Ssa1 S151D (S151D) were grown to exponential phase. Ssa1 was isolated by immunoprecipitation and binding partners were analyzed by label-free quantitative LC-MS/MS from 3 biological replicates. (A-C) Volcano plot comparison of Ssa1 binding partners between WT and SA (A), WT and SD (B), and SA and SD (C) summarizing results of t tests by showing log2 ratios of the fold change between comparisons (X axis) and the −log10 of p values (Y axis) for each binding partner identified. Values were normalized by levels of Ssa1 recovered from each immunoprecipitation. Error bars represent standard deviation. * indicates p < 0.05 by student 2-tailed T test. (D) Quantification of Ydj1 binding to WT, SA, or SD forms of Ssa1 in immunoprecipitations normalized by total lysates (“IP/Lysate”) compared to the levels in total lysates (“Lysate”). (E) Quantification of Sis1 binding to WT, SA, or SD forms of Ssa1 in immunoprecipitations normalized by total lysates (“IP/Lysate”) compared to the levels in total lysates (“Lysate”). Error bars represent standard deviation. * and ** indicate p < 0.05 and p<0.01, respectively, by student 2-tailed T test. (F) Δ*ssa1-4* yeast cells expressing Flag-Ssa1 (WT), Flag-Ssa1 S151A (S151A), or Flag-Ssa1 S151D (S151D) as well as galactose-inducible Ydj1 (Gal-Ydj1) or Sis1 (Gal-Sis1) were spotted on glucose or galactose-containing plates in fivefold serial dilution and grown for 3 days.

J-domain containing proteins (Hsp40s) and nucleotide-exchange factors (NEFs) are the major regulators of Hsp70 in its catalytic cycle (Saibil, 2013). Both co-chaperones dynamically interact with Hsp70 and carry out diverse functions. Hsp40 proteins transfer substrates to Hsp70 and promote ATP hydrolysis by Hsp70, which transforms Hsp70 into an ADP-bound closed conformation in which the substrate is tightly bound (Mayer, 2013). To complete the Hsp70 conformational cycle, NEFs promote the exchange of ADP for ATP and transform Hsp70 back to an ATP-bound open conformation which releases the folded client (Sharma *et al*, 2010). Two of the primary Hsp40 enzymes in S. *cerevisiae* are Ydj1 (Type I) and Sis1 (Type II), each of which independently direct Hsp70 to execute different cellular functions (Walters *et al*, 2015; Fan *et al*, 2004; Rüdiger *et al*, 2001). In our co-immunoprecipitation analysis, both Ydj1 and Sis1 are detected with wild-type Ssa1 and the S151A mutant, but show significantly lower association with Ssa1 S151D (Fig. 5D and 5E). In addition, we found that several known Hsp70 binding partners including endoplasmic reticulum (ER)-specific co-chaperone (HLJ1), mitochondria-specific ATPase (MCX1), and prion-associated factors (SUP35 and RNQ1) exhibited higher association with Ssa1 S151A compared to Ssa1 S151D (Fig. 5C, and Fig. S3). In contrast, several ribosomal proteins (both small and large subunit) exhibited higher association with Ssa1 S151D compared to Ssa1 S151A (Fig. 5C, Fig. S2, Table S3).

The binding defect between Ssa1 S151D and the Hsp40 factors could underlie the growth and heat survival deficits observed with this mutant. One possibility we considered was that the S151D phenotype might be suppressed by overexpression of the Hsp40 factors that exhibit lower levels of binding. To test this, we overexpressed Ydj1 and Sis1 with an inducible GAL promoter in cells expressing wild-type, S151A, or S151D Ssa1 genes at 30°C (Fig. 5F). Instead of rescuing the S151D growth defect, we found that the S151D-expressing cells were nearly inviable with galactose induction of Ydj1 and Sis1. Thus the reduction in Ydj1 and Sis1 binding cannot be functionally overcome by overexpression.

In addition to facilitating client binding by Hsp70 proteins, Ydj1 also promotes association between Ssa1 and Hsp104 as well as with small chaperones such as Hsp12, Hsp26, and Hsp31 to promote disaggregation and refolding of aggregated proteins (Fan *et al*, 2003; Cashikar *et al*, 2005; Duennwald *et al*, 2012). Perhaps relevant to this function, we observed that Hsp104 and small chaperones have higher protein expression in cells expressing Ssa1 S151D (Fig. S1). HSP82 (yeast HSP90), known to bind directly to Ssa1, is also present at higher levels in S151D-expressing cells (Fig. S1) (Kravats *et al*, 2018). Combined with the observation that HSF1 is hyperactive in cells expressing Ssa1 S151D, these observations suggest that Ssa1 phosphorylation may have broad effects on many clients through altered co-chaperone binding properties.

### Ssa1 S151 phosphorylation regulates disaggregation by Ssa1, Ydj1, and Hsp104

Previous studies have demonstrated that Ssa1 together with the Hsp40 co-chaperone Ydj1 are able to refold misfolded proteins and to prevent the formation of large aggregates from misfolded proteins (Balchin *et al*, 2016; Becker *et al*, 1996). Budding yeast also have the Hsp104 chaperone, which cooperates with Hsp70 and Hsp40 proteins to generate an efficient protein disaggregation assembly (Glover & Lindquist, 1998, 104). Hsp104 is a critical protein disaggregase for yeast cell survival of severe stress conditions, and has been shown to be responsible for extracting polypeptides from protein aggregates in cooperation with Ssa1 and Ydj1 (Lee *et al*, 2013).

To directly measure the effects of Ssa1 on protein folding, we established an in vitro luciferase-refolding assay with purified recombinant components (Fig. S4). We tested purified wild-type Ssa1, Ssa1 S151A, and Ssa1 S151D proteins in vitro with urea-denatured luciferase and found that purified Ssa1 S151D has poor luciferase reactivation efficiency compared to wild-type Ssa1 whereas the S151A mutant is more active than wild-type protein (Fig. 6A). In this case the Ssa1 proteins were produced in insect cells and the wild-type protein does have S151 phosphorylation (Fig. S4D) so it is expected that the wild-type would show an intermediate level of activity compared to the S151A and S151D mutants.

**Figure 6.**
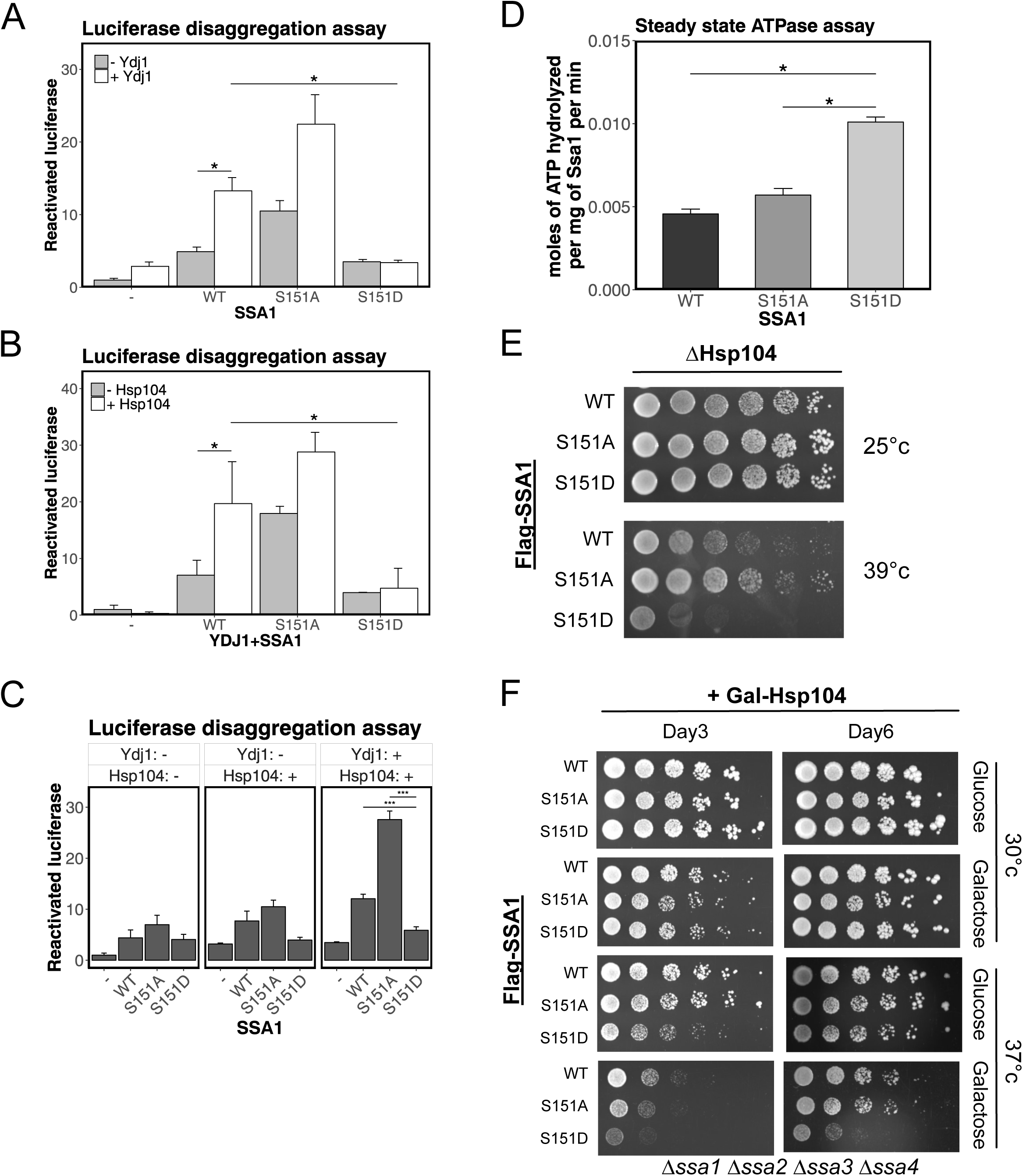
Ssa1 S151 phosphorylation regulates disaggregation by Ssa1, Ydj1, and Hsp104 *in vitro*. (A) Recombinant wild-type Ssa1, Ssa1 S151A, and Ssa1 S151D proteins (2.5 μM) were incubated with denatured luciferase (3.3 × 10^4^ units) in the presence or absence of Ydj1 (0.4 μM) and steady-state luciferase activity was measured. 3 biological replicates were performed and error bars represent standard deviation. * indicates p < 0.05 by student’s 2-tailed T test. (B) Recombinant wild-type Ssa1, Ssa1 S151A, Ssa1 S151D, and Ydj1 were tested for luciferase reactivation as in (A) but also in the presence of Hsp104 (1 μM) as indicated. (C) Comparative summary of luciferase assay results from (A) and (B). (D) Steady-state levels of ATP hydrolysis activity of recombinant wild-type Ssa1 (WT), Ssa1 S151A (S151A), and Ssa1 S151D (S151D). 3 technical replicates were performed and error bars represent standard deviation. * indicates p < 0.05 by student’s 2-tailed T test. (E) Δ*ssa1-4 Hsp104*Δ yeast cells expressing Flag-Ssa1 (WT), Flag-Ssa1 S151A (S151A), or Flag-Ssa1 S151D (S151D) were spotted in fivefold serial dilutions and exposed to 30 or 39°C for 120 hr. (F) Δ*ssa1-4* yeast cells expressing Flag-Ssa1 (WT), Flag-Ssa1 S151A (S151A), or Flag-Ssa1 S151D (S151D) as well as galactose-inducible Hsp104 (Gal-Hsp104) were spotted in fivefold serial dilutions and grown on glucose (Glu) or galactose (Gal) containing plates which were incubated at 30°C or 37°C for 3 days or 6 days as indicated.

Based on our co-immunoprecipitation observations, Ssa1 S151A and Ssa1 S151D mutants associate with Ydj1 differently, so Ydj1 might play a critical role in controlling refolding or disaggregation functions of Ssa1 in a manner that is controlled by S151 phosphorylation. We found that the combination of purified wild-type Ssa1 and Ydj1 was more efficient in reactivation of urea-denatured luciferase than reactions containing purified wild-type Ssa1 or purified Ydj1 only (Fig. 6A), consistent with previous reports (Glover & Lindquist, 1998, 104). Both wild-type and S151A proteins were significantly more efficient in luciferase reactivation with Ydj1 present while the activity of the S151D protein did not improve at all with Ydj1 addition (Fig. 6A), consistent with our binding results in Fig. 5 showing Ssa1 S151D has extremely reduced binding affinity for Ydj1.

Although Ssa1 and Ydj1 are able to perform a modest level of refolding, previous work has shown that Hsp104 can increase the level of luciferase reactivation by Ssa1 and Ydj1 (Glover & Lindquist, 1998). With purified Ssa1 and Ydj1 in the reaction, we thus compared the efficiency of the reaction with recombinant Hsp104 present (Fig. 6B). Consistent with previous results, we also observed that the addition of Hsp104 to wild-type Ssa1 and Ydj1 generated significantly higher levels of reactivated luciferase compared to reactions without Hsp104 (Fig. 6B). This cooperative effect was also observed with Ssa1 S151A and Ydj1, but Ssa1 S151D and Ydj1 fail to cooperate with Hsp104 in reactivation of aggregated luciferase (Fig. 6B). Without purified Ydj1, we observed that Ssa1 and Hsp104 failed to increase the level of luciferase reactivation although wild-type Ssa1 and Ssa1 S151A still showed more efficient luciferase reactivation than Ssa1 S151D (Fig. 6C). The results indicate that purified Ssa1 S151D fails to form an efficient and functional disaggregase complex with Hsp104 in vitro, at least in part due to lack of productive binding to Ydj1. This deficiency is not due to a lack of ATPase activity, as measurements of ATP hydrolysis with purified S151D protein show approximately 2-fold higher rates of hydrolysis compared with wild-type or S151A protein in vitro (Fig. 6D).

To determine if the effects of Ssa1 S151 phosphorylation are dependent on HSP104-dependent activities in vivo, we deleted the *HSP104* gene in our *Δssa1/2/3/4* yeast strain complemented by wild-type, S151A, or S151D *SSA1*. The cells were analyzed by serial dilutions on solid media and also exposed to heat shock at 39°C. The results show that Ssa1 S151D cells exhibit slow growth at elevated temperature compared to wild-type Ssa1-expressing cells while the Ssa1 S151A-expressing cells are even more resistant than wild-type cells (Fig. 6E). Thus, the phosphorylation of S151 (as it occurs in the wild-type strain) yields greater functional deficiencies in the absence of HSP104 than in its presence.

To test whether additional Hsp104 can recover the heat sensitivity of the S151D mutant strain, we introduced galactose-inducible Hsp104 into the *Δssa1/2/3/4* strain. Our results showed that additional Hsp104 expression did not affect growth of wild-type, Ssa1 S151A, or Ssa1 S151D cells at 30°C (Fig 6F). Interestingly, Ssa1 S151D cells with endogenous levels of Hsp104 (glucose) were sensitive to heat shock at 37°C with a short-term incubation (3 days), although recovered similar to wild-type cells after a long-term incubation (6 days). However, Ssa1 S151D cells with additional Hsp104 expression in the presence of galactose lost the ability to recover (Fig. 6F). The result indicates that additional Hsp104 might create incomplete or dominant negative disaggregase complexes in the presence of Ssa1 S151D that not only are non-functional but can block protein refolding.

### S151 phosphorylation regulates Sup35 prion function

In our co-immunoprecipitation analysis, we observed that Ssa1 is associated with two prion-associated proteins, Rnq1 and Sup35, and that the S151D form of Ssa1 exhibits significantly lower binding to these proteins (Fig. 5, S3). The *S. cerevisiae* [*PSI^+^*] prion is an inheritable, amyloid form of the Sup35 translation termination factor that is deficient in termination function (Liebman & Chernoff, 2012). Formation of the amyloid form occurs spontaneously but is promoted by Hsp70 function, specifically Ssa1, as well by the Hsp104 disaggregation machinery, which is required to convert large prion assemblies into smaller units that “seed” new fibers (Needham & Masison, 2008; Hung & Masison, 2006; Chernoff *et al*, 1995; Sharma & Masison, 2009; Shorter & Lindquist, 2004). Rnq1 has a prion-like domain that can functionally substitute for Sup35 and is required for the de novo appearance of [*PSI^+^*](Sondheimer & Lindquist, 2000; Derkatch *et al*, 2001). To test whether Ssa1 phosphorylation plays a role in prion propagation, we monitored [*PSI*^+^] using *Δssa1/2/3/4* yeast strain containing an *ade2-1* mutation as a color-based reporter for nonsense codon read-through (Fig. 7A)(Jung *et al*, 2000). In short, [*PSI*^+^] propagation leads to the generation of functional ADE2 protein (white) due to partial loss of Sup35 termination activity, whereas [*psi^−^*] cells have normal Sup35 activity and have a red pigment due to lack of ADE2 function. Expression of wild-type, S151A, and S151D versions of *SSA1* in a [*PSI*^+^] strain showed that [*PSI*^+^] propagation is maximal (white colonies) with S151A, while it is slightly less efficient in wild-type SSA1-expressing cells (slightly pink colonies) after long-term incubation (5 days)(Fig. 7B). In contrast, cells expressing Ssa1 S151D show no apparent [*PSI*^+^] activity (red colonies)(Fig. 7B). These results are consistent with our finding that Ssa1 S151A exhibits higher disaggregation efficiency in vitro whereas Ssa1 S151D fails to promote disaggregation under these conditions.

**Figure 7.**
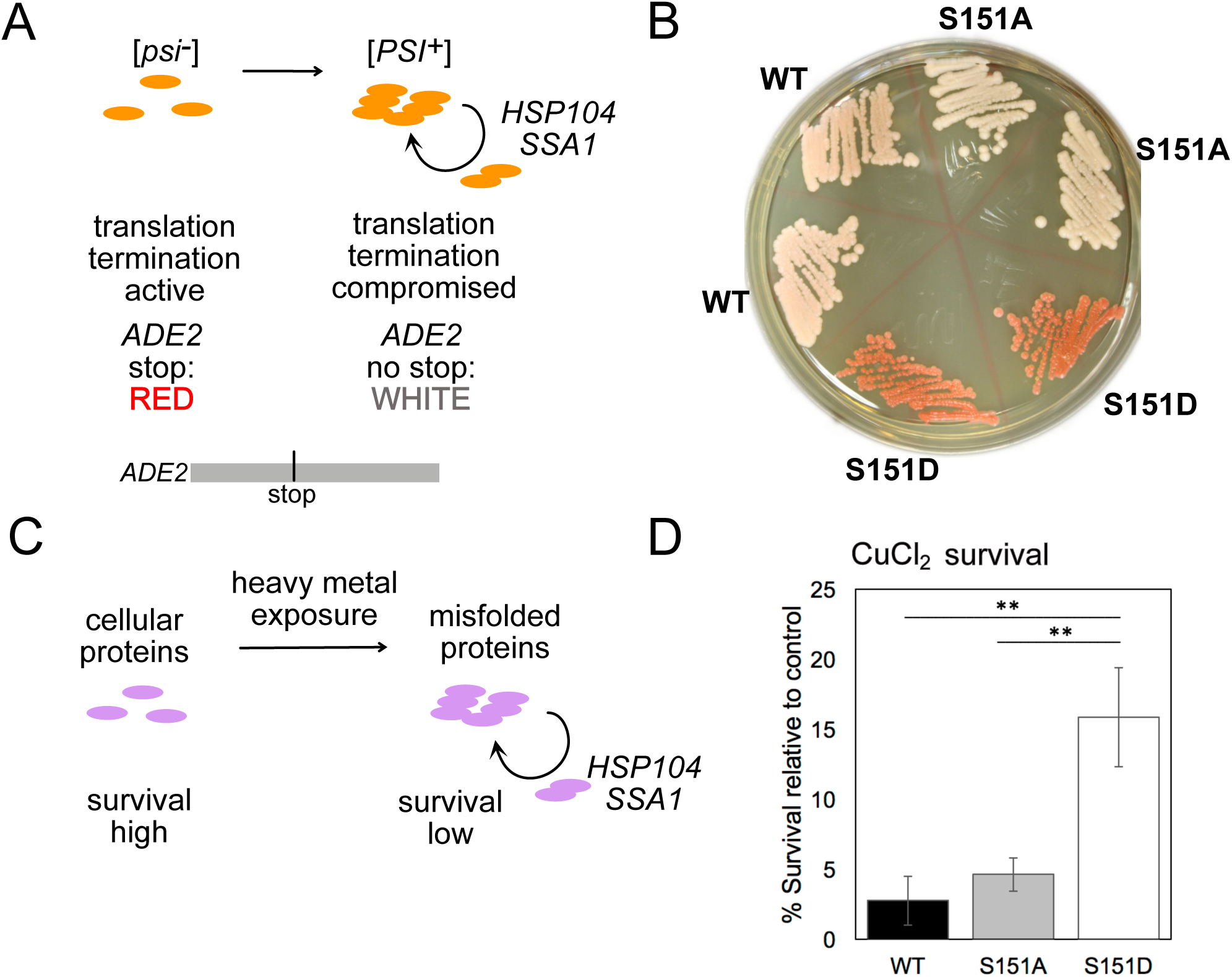
Prion propagation and heavy metal sensitivity are regulated by S151 Ssa1 phosphorylation. (A) Schematic diagram of *[PSI+]* Sup35 prion formation and propagation through Ssa1/Hsp104-dependent generation of seeds that form new aggregates. Red or white colonies are formed depending on the level of Sup35 function, as indicated. (B) *[PSI+]* Δ*ssa1-4* yeast cells expressing full-length Ssa1 WT, S151A, or S151D from a CEN plasmid were streaked on YPD plates for 5 days to show the level of *ade2-1* nonsense read-through. (C) Schematic diagram of heavy metal-induced protein aggregation and propagation through Ssa1/Hsp104-dependent generation of seeds that form new aggregates. (D) Δ*ssa1-4* yeast cells expressing full-length Ssa1 WT, S151A, or S151D from a CEN plasmid were exposed to CuCl_2_ (11 mM) for 6 hrs, followed by washing out of copper and plating on non-selective plates to determine viability. Survival relative to controls are shown for each strain. ** indicates t test p value < 0.01.

### S151 phosphorylation regulates survival of heavy metal exposure

An early report of Δ*hsp104* phenotypes by Lindquist and colleagues showed that cells lacking this chaperone are dramatically resistant to heavy metal exposure (cadmium and copper) compared to wild-type cells (Sanchez *et al*, 1992). More recent work suggests that cadmium and copper compounds directly generate misfolding of nascent proteins in budding yeast and higher organisms, and that these and other heavy metals generate metal-protein aggregates that seed the formation of new aggregates (Jacobson *et al*, 2017; Tamás *et al*, 2014; Hane & Leonenko, 2014). In this sense, metal-induced misfolding intermediates are analogous to prion intermediates in their ability to communicate protein misfolding states (Fig. 7C). To test if Ssa1 S153 phosphorylation may have a similar effect as Δ*hsp104*, we exposed yeast cells expressing wild-type, S151A, or S151D Ssa1 to copper (II) chloride and measured viability. It is clear that the phospho-mimic S151D allele promotes survival of copper exposure under these conditions at a level significantly higher than either S151A or wild-type Ssa1 expression (Fig. 7D), similar to the report for Δ*hsp104* (Sanchez *et al*, 1992).

### S151 phosphorylation regulates chaperone function in mammalian cells

As discussed above, Ssa1 S151 is highly conserved in eukaryotes including humans, where S153 is the corresponding residue in the constitutive Hsc70 as well as heat-induced HSP70 (Fig. 1A). Hsc70/Hsp70 phosphorylation at S153 was observed previously in a study of global SQ/TQ phosphorylation sites in human cells (Matsuoka *et al*, 2007), and we found this phosphorylation site in human U2OS osteosarcoma cells as well (Table S2). To confirm that phosphorylation occurs during normal growth, we expressed V5-tagged wild-type HSC70 and a S153A mutant in U2OS cells, isolated the protein by immunoprecipitation, followed by western blotting with the phospho-specific antibody. The results confirm that S153 phosphorylation does occur in these cells, although residual signal is still present with the S153A mutant, perhaps due to cross-reacting phosphorylation elsewhere in the protein (Fig. 8A).

**Figure 8.**
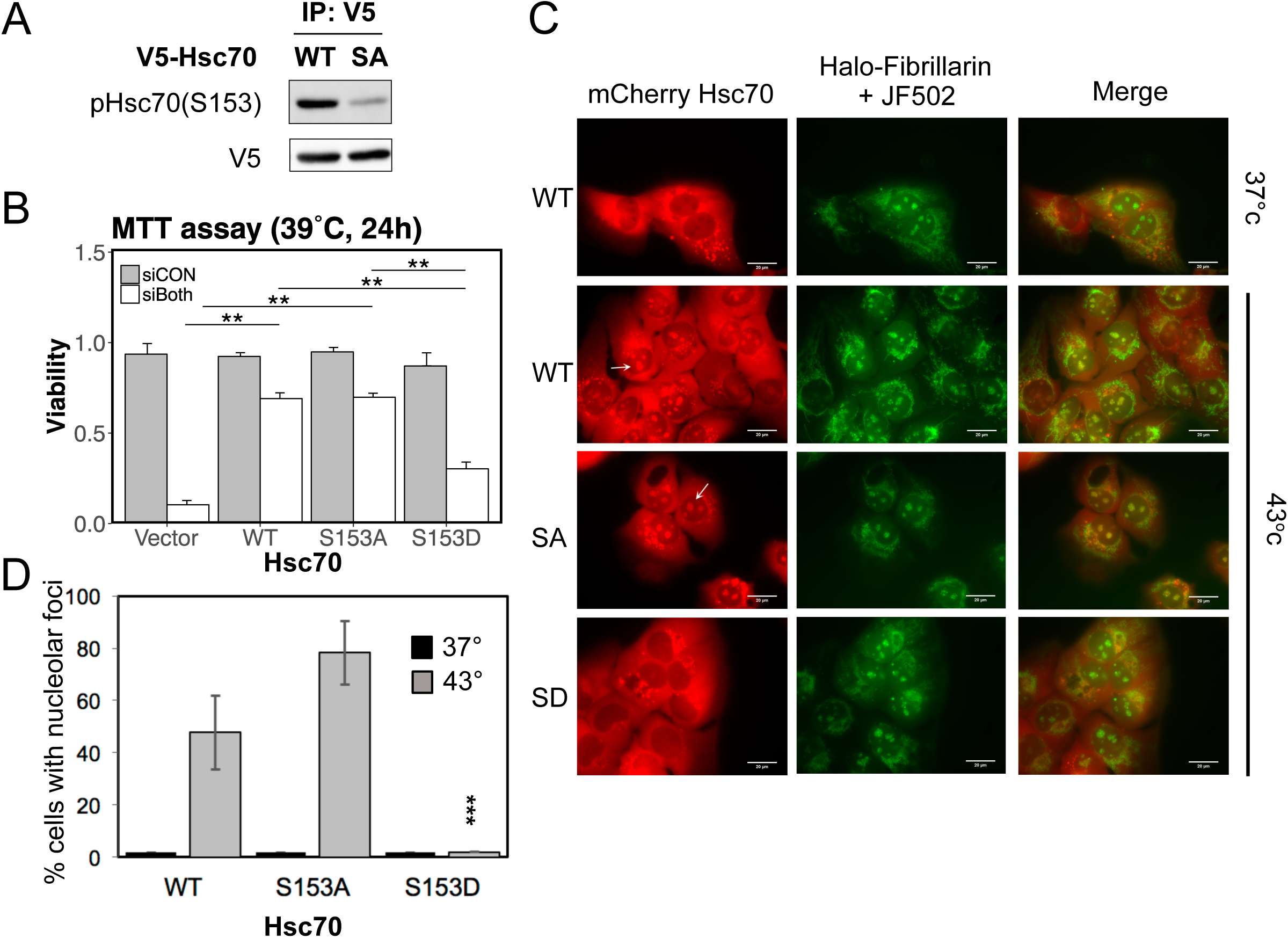
S151 phosphorylation occurs in mammalian cells and regulates heat-induces relocalization of HSC70. (A) V5-Hsc70 (WT) or V5-Hsc70 S153A (SA) were expressed in U2OS cells with concurrent Hsc70/Hsp70 depletion. Hsc70 was isolated by immunoprecipitation and analyzed by Western blotting with anti-phospho-Ssa1 S151 and anti-V5 antibodies. (B) U2OS cells expressing V5-Hsc70 (WT), V5-Hsc70 S153A (S153A), or V5-Hsc70 S153D (S153D) were transfected with control siRNA (siCON) or siRNAs directed against HSPA8(HSC70) and HSPA1A(HSP70)(siBoth) and seeded in 96-wells plates. Cells were treated with doxycylin to induce recombinant Hsc70 expression and incubated at 39°C for 24 hr. Cell viability was measured by MTT assay (Thermo Fisher Scientific). 3 biological replicates were performed and error bars represent standard deviation. * indicates p < 0.05 by student’s 2-tailed T test. (C) mCherry-tagged WT, S153A, or S153D Hsc70 was expressed in U2OS cells expressing Halo-Fibrillarin and also treated with the JF502 halotag ligand. Cells were exposed to heat shock (43°C) for 60 min and analyzed by fluorescence microscopy. White arrows indicate nucleolar Hsc70. (D) Quantification of results from (C) showing percentage of cells with overlap between mCherry-Hsc70 and Halo-Fibrillarin. *** indicates t test p value < 0.0001 comparing S153D to either wild-type or S153A.

To investigate the role of Hsc70 phosphorylation at S153 we depleted endogenous Hsc70 using siRNA; however, in human cells, depletion of Hsc70 generates a dramatic induction of Hsp70 (HSPA1A/B) expression (Fig. S5) (Powers *et al*, 2008). To alleviate this overexpression we also depleted Hsp70, as previously described (Powers *et al*, 2008), resulting in 4-fold lower levels of Hsc70 with approximately 3-fold higher levels of Hsp70 relative to untreated cells. In these double-depleted cells we expressed V5-tagged wild-type Hsc70, the non-phosphorylatable mutant Hsc70 (S153A) or the phospho-mimetic mutant Hsc70 (S151D) from a stably-integrated doxycyline-inducible promoter (Fig. S5).

We tested for the effect of S153 phosphorylation status on survival of 39°C heat exposure and observed that cells expressing the phospho-mimetic S153D mutant were hypersensitive to heat shock (Fig. 8B), consistent with the results we observed in yeast cells (Fig. 1). Therefore, Hsc70 S153 phosphorylation also acts in a dominant negative way during heat shock in human cells.

Hsc70/Hsp70 proteins are critical in the nucleolus for ribosome biogenesis, stress responses, and cell signaling (Bański *et al*, 2010). In mammalian cells during heat shock, Hsc70 rapidly accumulates in nucleoli, a response which is important for counteracting damage and protein misfolding during heat stress (Kose *et al*, 2012; Frottin *et al*, 2019). We tested nucleoli accumulation of GFP-tagged wild-type, S153A, or S153D Hsc70 in U2OS cells during heat shock and found that Hsc70 S153D completely failed to accumulate in nucleoli compared to wild-type and Hsc70 S153A cells (Fig. 8C, D). Thus, S153 phosphorylation regulates Hsc70 relocalization in response to heat in a similar manner in humans as it does in budding yeast.

## Discussion

Here we investigated the role of Ssa1/HSC70 phosphorylation at S151(S153) during normal growth as well as stress conditions. We found that Ssa1 S151 phosphorylation is promoted by Cdk1 activity during logarithmic growth, and that this modification negatively regulates survival of heat exposure as well as starvation conditions. Ssa1 phosphorylation at S151 alters the interaction with Hsp40 co-chaperones that further impact the formation and function of Hsp104-dependent disaggregase complexes. Localization patterns of Ssa1 and Hsc70 in response to heat stress are also very sensitive to S151(S153) modification, as is survival of heavy metals and the maintenance of the [*PSI+*] prion in budding yeast. Thus Cdk1 appears to control many aspects of Ssa1/HSC70 function, and may regulate the ability of prions to affect gene expression.

### The effect of chaperone phosphorylation on Ssa1 function

Hsp70 includes 3 major domains: the N-terminal nucleotide binding domain (NBD), a linker region, and the C-terminal substrate binding domain (SBD)(Mayer, 2013). The ATPase activity of Hsp70 is mainly dependent on the NBD, a globular structure composed of four coordinated subdomains, each of which is responsible for specific interactions with ATP. However, previous studies of bacterial DnaK and DnaJ show that NBD ATPase activity is also tightly coupled to SBD interactions as well as to the binding of co-chaperone and peptide clients (Flynn *et al*, 1989; Buchberger *et al*, 1995; Bukau & Horwich, 1998). Based on DnaK strutures as well as the Hsp70 ortholog Sse1, serine 151 of Ssa1 is predicted to be in a surface-exposed loop of the NBD, juxtaposed against the SBD (Kityk *et al*, 2012; Liu & Hendrickson, 2007). Taken together, this data suggests that S151 phosphorylation may not only play a regulatory role in domain interaction between NBD and SBD, but also in co-chaperone interactions.

Consistent with this prediction, we found that the phospho-mimetic version of S151, Ssa1 S151D, exhibits very low levels of binding to Ydj1 and Sis1: co-chaperones in the Hsp40 family responsible for escorting clients to Hsp70 and accelerating HSP70 ATP hydrolysis (Cheetham & Caplan, 1998; Greene *et al*, 1998; Jiang *et al*, 2007). Results with recombinant Ssa1 and Ydj1 in an *in vitro* luciferase refolding assay showed that non-phosphorylated Ssa1 (the S151A phospho-blocking mutant) exhibits even higher refolding efficiency than the wild-type Ssa1 protein and that addition of Ydj1 cooperatively increases this activity, as reported previously for Ssa1/Ydj1 (Glover & Lindquist, 1998). In contrast, the S151D mutant failed to promote any co-chaperone-mediated refolding, and expression of the Ssa1 S151D phospho-mimic allele *in vivo* results in accumulation of higher levels of more protein aggregates in vivo (Fig. 2).

The Hsp104 chaperone is an important disaggregase in budding yeast, functioning cooperatively with Hsp70 and Hsp40 proteins to recognize aggregated proteins and extract individual polypeptides (Glover & Lindquist, 1998; Lee *et al*, 2013). Here we found that the S151D phospho-mimic Ssa1 failed to cooperate with Hsp104 in luciferase refolding assays in vitro and also exhibited more severe heat shock sensitivity in a *Δhsp104* background in vivo. Under these conditions, the S151A form of Ssa1 was remarkably more efficient than wild-type Ssa1 in promoting heat shock survival, suggesting that the unphosphorylated form of Ssa1 can partially compensate for chaperone function normally provided by Hsp104. Expressing higher levels of Hsp104, Ydj1, and Sis1 in Ssa1 S151D strains failed to improve heat shock survival, thus the functional defect with S151D Ssa1 is not simply a lower affinity of Ssa1 for the co-chaperones but more likely involves a conformational change that is incompatible with co-chaperone function. A similar combination of attributes was reported with Cdk1 phosphorylation of Ssa1 on T36, also in the NBD (Truman *et al*, 2012). In this case, the phospho-mimic version of Ssa1 (T36E) exhibited higher levels of nucleotide binding *in vitro*, low survival of heat shock, reduced binding to Ydj1 (although in this case no effect on Sis1 binding), and altered cell cycle progression (Truman *et al*, 2012). The T36 residue is adjacent to the ATP-binding pocket in Ssa1, unlike S151, probably explaining the change in nucleotide binding affinity, but similarly illustrates dominant negative effects of Cdk1 phosphorylation on Ssa1 functional interactions.

### Ssa1 S151 phosphorylation occurs under conditions of rapid growth

Although Ssa1 S151 does not conform to the S/T-P consensus phosphorylation motif that is normally found for Cdk1 substrates, our data from *in vitro* as well as *in vivo* assays indicates that Cdk1 participates in Ssa1 phosphorylation at S151 (Fig. 3). Non-S/T-P sites for Cdk kinases have been reported in other biological contexts (Suzuki *et al*, 2015; Kusubata *et al*, 1993; Satterwhite, 1992; Harvey *et al*, 2005; Isoda *et al*, 2011; McCusker *et al*, 2007). Ssa1 S151 was also identified in a phosphorylation screen for Cdk1 targets, although the extent of cell cycle dependence was not very high compared to other targets, suggesting that there could also be other kinases responsible for this modification (Holt *et al*, 2009). Our results are not inconsistent with this conclusion as we do not see an absolute dependence on Cdk1 *in vivo*. We do observe a strong reduction in S151 phosphorylation with Tor inhibition and in stationary phase (Fig. 3), suggesting that if there are other kinases involved they are likely also subject to growth regulation.

Consistent with the idea that S151 is generally unphosphorylated during conditions of growth inhibition, we found that cells expressing Ssa1 S151A exhibit significantly better growth in the presence of the Tor inhibitor rapamycin compared to cells expressing either wild-type or S151D Ssa1 (Fig. 3). Previous proteomic analysis of yeast cells grown in the presence of rapamycin indicated that cells with activated Hsf1 are hypersensitive to rapamycin (Bandhakavi *et al*, 2008). Our finding that Ssa1 S151D-expressing cells show high levels of Hsf1 activation (Fig. 1) is consistent with this, and suggests that regulation of S151 modifications contribute to survival during nutrient limitation. Ssa1 S151D also exhibits lower levels of binding to ribosomal subunits (Fig. S2), a deficit that appears not to be growth-limiting under normal conditions but may be important for viability during starvation.

### Negative functional effects of S151 phosphorylation

The S151D phospho-mimic allele of Ssa1, as well as the S153D form of human HSC70, have mostly negative-acting functional effects on survival of stress conditions and growth. The S151A allele, on the other hand, exhibits either similar or higher activity than the wild-type S151 in these assays, with higher activity associated with Tor inhibition or Hsp104 deficiency. Most prokaryotes have an alanine at this position, so it is puzzling that eukaryotes have stably inherited and maintained a version of the chaperone that can be inactivated by a kinase that is active during normal growth.

One possibility is that the multiplication of Hsp70 orthologs in eukaryotes has selected for a diversification of functions and binding partners (Albanèse *et al*, 2006). In *S. cerevisiae*, the *SSA* subfamily (Ssa1, Ssa2, Ssa3, Ssa4) is important for protein folding, membrane translocation, nuclear import, and transcriptional responses to a variety of stress conditions, while the *SSB* subfamily (Ssb1 and Ssb2) are key components of the ribosome-associated complex (RAC) that assists the de novo folding of newly synthesized polypeptides (Boorstein *et al*, 1994; Kim *et al*, 1998; Oka *et al*, 1998; Shulga *et al*, 1996; Mayer & Bukau, 1998; Stone & Craig, 1990; Jaiswal *et al*, 2011). Ssb1 and Ssb2 have an alanine at the 151 position, while all 4 of the Ssa proteins have a serine. It could be that the ability to phosphorylate Ssa proteins, which is predicted to block Ydj1 and Sis1 binding, could promote associations that are beneficial in other environmental conditions. These alterations are likely tolerated because of the presence of the ubiquitiously expressed, non-phosphorylatable Ssb proteins. Although most prokaryotes have an alanine at the equivalent position and most eukaryotes have a serine, the NCBI database shows approximately 120 bacterial and ascomycete species with an aspartate. How the Hsp70 proteins with this sequence function in concert with their respective DnaJ(Hsp40) co-chaperones is unknown.

### Prions and metal-induced misfolded proteins as targets of Hsp70 activity

We show in this work that the [*PSI^+^*] prion can be regulated by modification of S151 of Ssa1. Hsp70 is well-known for its role in prion dynamics, as many laboratories have documented the necessity of Hsp70, Hsp40, and Hsp104 protein families for propagation of the amyloid structures that constitute the infectious and heritable species (Liebman & Chernoff, 2012; Chernoff *et al*, 1995; Jung *et al*, 2000; Shorter & Lindquist, 2005). Previous studies of Hsp70 mutants showed that an L483W change in Ssa1 generates higher rates of ATP hydrolysis, reduced co-chaperone binding, and reduced protein refolding efficiency in comparison to wild-type Ssa1, as well as dramatically reduced ability of the protein to promote Sup35-dependent prion propagation (Needham & Masison, 2008; Jones & Masison, 2003), all very similar to the Ssa1 S151D mutant described here.

The idea that misfolded proteins can form seeds that spread misfolding to other, non aggregated protein species is common to both prions as well as to nascent proteins exposed to heavy metals (Jacobson *et al*, 2017; Tamás *et al*, 2014; Hane & Leonenko, 2014). In this sense, optimal chaperone function could be non-productive as it can generate new seeds from aggregated species and produce misfolded protein complexes at much higher rates compared to rates in the absence of chaperones. The higher viability conferred by S151D Ssa1 in the presence of copper shown here suggests that there could be selection for phosphorylation due to pervasive heavy metals in the environment (Tchounwou *et al*, 2012) that directly induce protein misfolding.

Our finding that Cdk1 is involved in phosphorylation of S151 suggests that [*PSI^+^*] prion as well as metal-induced misfolded protein species would tend to be repressed by phospho(S151)-Ssa1 in actively growing cultures. [*PSI^+^*] and other related prions in yeast affect translation read-through - a phenomenon proposed to increase the diversity of expressed proteins by translation of 3ʹ UTR regions and other normally untranslated sequences (Shorter & Lindquist, 2005). It is attractive to consider the possibility that serine 151 phosphorylation is a mechanism by which this evolutionary diversification could be controlled, in effect a switch regulating the appearance of novel polypeptides that is dependent on stress conditions and growth rate. The evolutionary maintenance of S151 in eukaryotic Hsp70 orthologs, perhaps driven by metal exposure, could ultimately regulate diversification advantages through prion regulation of gene expression in a fluctuating environment.

## Materials and Methods

### Yeast strains and plasmids

Yeast strains and plasmids used in this study are listed in Table S4. *S. cerevisiae* yeast cultures were grown in synthetic minimal defined media (0.67% yeast nitrogen base without amino acids, ammonium sulfate, and appropriate amino acids) with 2% glucose or YPAD media (1% yeast extract, 2% Becto peptone, 0.004% Adenine Hemisulfate) with 2% glucose. pESC-URA-GFP-Ubc9ts was a gift from Judith Frydman (Addgene plasmid # 20369; http://n2t.net/addgene:20369; RRID:Addgene_20369) (Kaganovich *et al*, 2008). HSP104-b/Leu(WT) was a gift from Susan Lindquist (Addgene plasmid # 1156; http://n2t.net/addgene:1156; RRID:Addgene_1156) (Schirmer *et al*, 2004). 5787 pET28aSX104B/ pES42 was a gift from Susan Lindquist (Addgene plasmid # 1229; http://n2t.net/addgene:1229; RRID:Addgene_1229) (Shorter & Lindquist, 2004). pRS315-SSA1 and pDP122 were gifts from Daniel Masison and David Pincus, respectively.

### Yeast protein extraction and western blotting

Yeast cells were grown in 2 mL culture media for 2 days to stationary phase and inoculated into larger volume cultures at 0.15 O.D. _600_ per ml. Cells were incubated at 30°C until log phase (0.3 – 1 O.D. _600_) and collected at 4000 rpm for 5 min. Protein was extracted by bead beating in 0.3 ml lysis buffer [25 mM Tris-HCl buffer pH 7.4, 150 mM NaCl, 1 mM EDTA, 10% Glycerol, 0.5% NP40, 1 mM DTT, EDTA-free protease inhibitor mini tablets (Pierce), EDTA-free phosphatase inhibitor mini tablets (Pierce)] and 0.1ml acid-washed glass beads for 1 min at room temperature. Protein lysates were collected after 3500 rpm for 5 min at 4°C. 400 μg of protein lysates incubated with 50 μl anti-Flag-magnetic beads (MBL) for 1 hour at 4°C. Beads were collected and washed 3 times with 500 μl lysis buffer. After the third wash, the lysis buffer was removed completely and the beads were mixed with 20 μl 2.5x SDS sample buffer. The mixture was boiled at 95°C for 5 min. Boiled samples were loaded into 8% SDS-PAGE or NuPAGE 4% - 12% Bis-Tris protein gels (ThermoFisher). Specific protein targets were analyzed by immunoblot assay with specific antibodies listed in Table. S1. The S151 phospho-specific antibody was produced by PhosphoSolutions using a peptide containing the phosphorylated site.

### Galactose induction

Yeast cells were grown in 2 ml culture media with 2% glucose overnight and then inoculated into 25 ml of synthetic minimal defined media containing 2% raffinose overnight. This culture was used to inoculate cultures at 0.15 O.D. _600_ 2% raffinose and incubated at 30°C until log phase (0.3 – 1 O.D. _600_). 3X YP media (3% Yeast extract, 6% Peptone with 6% Galactose) was used to induce GAL expression (or glucose-containing media as a control).

### Copper sensitivity

*Δssa1-4* yeast cells expressing wild-type, S151A, or S151D Ssa1 mutants were grown to log phase in synthetic minimal defined media (∼0.4 to 0.5 O.D. _600_); 11 mM CuCl_2_ was added for 6 hrs; cells were washed with YPAD media, and plated in dilutions on YPAD plates. Colony survival was measured with 3 biological replicate experiments comparing copper exposure to control cultures.

### Protein aggregate isolation and analysis

The assay was performed as described previously (Koplin *et al*, 2010). To prepare cell lysates, the pellets were resuspended in lysis buffer [20 mM Na-phosphate (pH 6.8), 10 mM DTT, 1 mM EDTA, 0.1% Tween 20, 1 mM PMSF, and EDTA-free protease inhibitor mini tablets (Pierce)]. Cells were lysed in a 4°C water bath–based sonicator (Bioruptor; eight times at level 4.5 and 50% duty cycle) and centrifuged for 20 min at 200xg at 4°C. Supernatants were adjusted to the same concentration, and protein aggregates were pelleted at 16,000x g for 20 min at 4°C. After removing supernatants, protein aggregates were washed twice with 2% NP-40 [20 mM Na-phosphate (pH 6.8), 1 mM PMSF, and EDTA-free protease inhibitor mini tablets (Pierce)], sonicated (six times at level 4.5 and 50% duty cycle), and centrifuged at 16,000 xg for 20 min at 4°C. Aggregated proteins were washed in buffer without NP-40 (sonication; four times at level 3 and 50% duty cycle), boiled in 2.5x SDS sample buffer, separated in NuPAGE 4% - 12% Bis-Tris protein gels (ThermoFisher), and analyzed by coomassie blue staining. Levels of aggregates were quantified using Image Studio.

### Filter aided sample preparation and trypsin digestion for mass spectrometry

Detergent-resistant aggregates or immunoprecipitated samples were resuspended in 15 μl of 10% SDS sample buffer and 50 mM beta-mercaptoethanol and boiled in 100°C for 5 minutes. The samples were diluted with 200 μl of UA buffer (8 M Urea, 0.1 M Tris-HCl pH 8.8) at room temperature. Microcon®-30 centrifugal filter units (Millipore, MRCF0R030) were equilibrated with 20% ACN, 2% formic acid solution (14,000 xg for 10 min) prior to use. Diluted samples were loaded on the filters then washed with UA buffer (8 M Urea, 0.1 M Tris-HCl pH 8.8) 3 times. After washing, samples were reduced with 50 mM DTT in UA buffer (8 M Urea, 0.1 M Tris-HCl pH 8.8) which was added to filters, incubated 5 minutes at room temperature and spun off. Then samples were alkylated with 50 mM iodoacetamide in UA buffer (8 M Urea, 0.1 M Tris-HCl pH 8.8), incubated 5 minutes at room temperature and spun off. Samples were de-salted with 40 mM ammonium bicarbonate (ABC) 3 times. 100 ul 40 mM ABC with 0.5 μl of trypsin gold (Promega, V528A) in PBS was added to samples and were incubated overnight (37°C). Trypsinized peptides were eluted by centrifugation; filters were washed with 20% ACN, 2% formic acid solution and filtrate was combined with trypsinized peptides eluted in ABC. Peptide samples were dried by lyophilization, de-salted with C18 tips (Pierce, QK224796) according to manufacturer’s instructions, and resuspended in 80% ACN 2% formic acid for LC-MS-MS analysis at the Proteomics Core Facility (University of Texas at Austin). All centrifugations were done at 14,000 xg for 20 min unless otherwise noted.

### Recombinant protein expression

Wild-type Ssa1, Ssa1 S151A, and Ssa1 S151D proteins were expressed using the Bac-to-Bac baculovirus system (ThermoFisher). SSA1 wild-type, SSA1 (S151A), and SSA1 (S151D) genes were cloned into pFastBac1, Flag-tagged at the N-terminus (generating pTP4416, pTP4417, and pTP4418, respectively). Recombinant bacmid DNA derived from these transfer vectors was transfected into Sf21 insect cells for recombinant baculovirus production according to manufacturer instructions. Ssa1 proteins were expressed in Sf21 insect cells after baculovirus infection. Cell pellets were lysed by homogenization and sonicated three times for 40 seconds in A buffer (25 mM Tris pH 7.4, 100 mM NaCl, 10% glycerol, and 2 mM dithiothreitol (DTT)) containing 0.5% tween-20, 1mM phenylmethylsulfonyl fluoride (PMSF), and 0.001 % 2-Mercaptoethanol. The lysate was centrifuged for 1 hr at 35000 rpm at 4°C. The supernatant was incubated with ∼1mL M2 anti-Flag antibody-conjugated agarose resin (Sigma) with rotation at 4°C for 1hr. After incubation the lysate with resin was centrifuged for 3 min at 1000Xg. After removing the supernatant, the remaining resin was washed with 20 mL of A buffer twice and was eluted with 5mL of A buffer containing 0.8 mg/mL 3X Flag peptide (Sigma). The peptide was incubated with the resin for 20 min before elution. The Flag elution was then loaded onto 1mL HiTrap Q column (G.E.) and washed with buffer A then eluted with buffer A containing 500 mM NaCl. The eluted protein fractions were dialyzed in A buffer and the dialyzed fractions were aliquoted frozen in liquid nitrogen, and stored at −80°C. Protein concentration was quantified by SDS-PAGE and coomassie blue staining using a Licor Odyssey Imager.

V5-tagged Ydj1 proteins were purified as GST fusion proteins in BL21 *E. coli*. Starter cultures were prepared at 37°C for 16 hr. Overnight cultures were diluted 1:20 for additional 2 hr incubation at 37°C. 100 μM Isopropyl b-D-1-thiogalactopyranoside (IPTG) was used to induce protein expression for 3 hr at 37°C. Cells pellets were collected using 3400 rpm for 10 min, resuspened in phosphate buffer saline (PBS) with 1% Triton, and sonicated for 30 seconds. Cell supernatants were collected using 10000 rpm for 10 min and incubated with Glutathione sepharose 4B resin (GE Healthcare) for 2 hr at 4°C. Beads were collected and washed three times with PBS 1% Triton and three times with 50mM Tris-HCl pH8.0, 150mM NaCl, 0.01% Triton, and 2.5mM EDTA. GST fusion V5-Ydj1 was subjected to site-specific cleavage with PreScission Protease (GE Healthcare) to remove the GST tag. V5-Ydj1 was purified from beads and its protein concentration was quantified by SDS-PAGE using a Licor Odyssey Imager.

Flag-Hsp104 was purified from BL21 *E. coli*. Starter cultures were prepared at 37°C for 16 hr. Overnight cultures were diluted 1:20 for additional 2 hr incubation at 37°C. 100 μM Isopropyl b-D-1-thiogalactopyranoside (IPTG) was used to induce protein expression for 3 hr at 37°C. Cells pellets were collected using 3400 rpm for 10 min. The protein purification protocol step follows as previously described in Ssa1 protein purification.

### Luciferase refolding assay

Firefly luciferase (Sigma L9420) was diluted in refolding buffer (20 mM Tris-HCl pH 7.4, 50 mM KCl, and 5 mM MgCl_2_) 5-fold and denatured with 6 M Urea for 30 min at room temperature. Denatured luciferase was then diluted 100 fold with refolding buffer and incubated with indicated chaperones, indicated co-chaperones, 10 mM ATP, and 10 mM DTT for a final volume of 25 μl. The reaction was incubated at 30°C for 30 min. 5 μl reaction were mixed with 50 μl luciferase assay substrate (Promega). A Tecan SPARK 10M plate reader was used to measure luciferase activity.

### ATP hydrolysis measurements

Recombinant proteins were prepared in Buffer A (25mM Tris-HCl pH 8.0, 100mM NaCl, 10% Glycerol, 1mM DTT). Reactions were started by adding [α-^32^P] ATP mixtures (25 mM Mops pH7.0, 5mM MgCl_2_, 0.2 mM DTT, 0.1 mg/mL BSA, 50μM ATP, 50nM [α-^32^P] ATP) to protein solutions, followed by incubation at 37°C for 1 hr. 1ul of Stop solution (2% SDS and 100 mM EDTA) was added to stop the reaction. 1 μl of each reaction was spotted onto a Plastic Backed TLC Cellulose PEI plate (Scientific Adsorbents Incorporated, #78601). The plate was dried and run in 0.75 mM KH_2_PO_4_ (monobasic) buffer. Percentage of hydrolyzed [α-^32^P] ATP and ADP were quantified by Typhoon FLA 7000 biomolecular image and normalized by the reaction without proteins. Normalized percentage of hydrolyzed ATP was divided by the amount of Ssa1 proteins and further divided by the incubation time of the assay.

### GFP-Ubc9^ts^ protein aggregate analysis

Galactose-controlled GFP-Ubc9^ts^ was used as an indicator of the level of protein aggregation. Cells were grown overnight and re-inoculated into 2 ml of synthetic minimal defined media with 2% raffinose overnight. Cells were inculated at 0.15 O.D. _600_ per ml in 2% raffinose and incubated at 30°C until log phase (0.3 – 1 O.D. _600_). Synthetic minimal defined media with 3xYP(2% Galactose) was added to induce expression of GFP-Ubc9^ts^ (or 3xYP with 2% Glucose as a control). 54°C culture media was prepared ahead and used to perform 42°C heat shock in a 1:1 ratio of culture media. Heated-cells were incubated at 42°C for 30min and then prepared for imaging. Cells with at least one focus was counted as positive and compared to the total number of cells containing GFP fluorescence.

### GFP-Ssa1 cellular foci analysis

For the starvation experiment, GPF-tagged wild-type Ssa1, S151A, and S151D cells were grown overnight and re-inoculated into 2 ml of synthetic minimal defined media with 2% glucose in a concentration of 0.15 O.D. 600 per ml. Cells were grown to log phase, collected by centrifugation, and then resuspended in media lacking glucose for 10 min before imaging. Cells with at least one focus was counted as positive and compared to the total number of cells containing GFP fluorescence. For the saturation phase experiment, GPF-tagged wild-type Ssa1, S151A, and S151D cells were grown for 2 days to saturation phase. Cells with at least one focus was counted as positive and compared to the total number of cells containing GFP fluorescence.

### HSE-YFP reporter heat shock assays

The HSE-YFP reporter, pNH605-4xHSEpr-YFP (pDP122, a gift from David Pincus), was integrated into the genome of the *ssa1-4*Δ yeast cells (a gift from Sabine Rospert). This strain was complemented with Flag-tagged Ssa1 (wild-type, S151A mutant, or S151D mutant). The assay was performed essentially as described (Zheng *et al*, 2016). Briefly, cells were prepared at a density of 0.2 O.D. 600 per ml and incubated at the indicated temperature on a thermal mixer. After 30 or 60 min, 50 μl were collected and treated with cycloheximide (Final concentration: 50 μg / mL) to stop translation. Additional 2 hr incubation at 30°C is required to mature YFP fluorophores. The Mean Fluorescence Intensity (MFI) of each cell was measured using a BD LSRFortesa™ flow cytometer and analyzed by FlowJo.

### Double siRNA knockdown in human cells

siRNA sequences were designed based on a previous study (Powers *et al*, 2008) and are listed in Table S4. The transfection of siRNA into U2OS cells was performd with Oligofectamine transfection reagent (ThermoFisher) according to manufacturer instructions. Double knockdowns were performed with 20 μM of each siRNA.

### Heat shock and nucleolin staining of human cells

U2OS cells expressing mCherry-Hsc70 (WT, S153A, S153D) and Halo-Fibrillarin were plated into WillCo-dish® Glass Bottom dishes (cat. 14023-200) with cell culture media containing 1ug/ml doxycycline a day before the experiment. Cells were heat shocked for 45 minutes in a 43°C tissue culture incubator (5% CO_2_). 1 pM JF502 halotag ligand (a gift from Luke Lavis) was added to cell media for 15 minutes. Finally cells were washed with sterile PBS in room temperature then analyzed using a Zeiss Axiovert Fluorescent Light Microscope with a 64x oil immersion objective. Images were analyzed with FIJI software (ImageJ v1.52c). For quantification, at least 70 cells from several fields of view were scored for GFP-HSC70 foci in the nucleolus (overlapping with Halo-Fibrillarin) under normal growth conditions (37° C) and heat shock (43° C).

### Isolation of tagged Hsc70 from human cells

Biotinylated, V5-tagged Hsc70 was expressed in human U2OS cells and isolated with streptavidin-coated beads under denaturing conditions. Briefly, cells were lysed with urea solution (8M Urea, 50mM Tris pH8, 5mM CaCl_2_, 30mM NaCl, 0.1% SDS and 1:1000 PMSF/Protease Inhibitor). Lysates were sonicated for 30 sec. 3 mg of lysates were diluted to 1M Urea. 120 μL of streptavidin magnetic beads were added to each samples and rotated overnight at RT. The next day, beads were washed twice for 30 minutes each with 1M Urea solution (1M Urea, 50mM Tris pH8, 5mM CaCl_2_, 30mM NaCl, 0.1% SDS) at RT. Then beads were washed with LiCl (500mM) for 30 minutes at RT, then washed 3 times with 0.1%SDS, 0.2%SDS and 0.5%SDS for 30 minutes at RT respectively. Finally, samples were eluted with 1% SDS with 2-mercaptoethanol at 100°C. Samples were frozen in −20°C until analyzed by filter-assisted sample preparation and trypsinization (see above).

## Acknowledgments

We are indebted to several laboratories for yeast strains and reagents: Sabine Rospert, David Pincus, David Morgan, Daniel Masison, and Luke Lavis, as well as help with expression constructs from Caleb Swaim. We also thank Judith Frydman for pESC-URA-GFP-Ubc9ts (Addgene plasmid # 20369) and Susan Lindquist for HSP104-b/Leu(WT)(Addgene plasmid # 1156) and pET28aSX104B/ pES42 (Addgene plasmid # 1229). We thank Daniel Masison and members of the Paull laboratory for helpful comments. C-H. Kao was supported in part by a Technologies Incubation Scholarship from the Taiwan Ministry of Education.

## References

Albanèse V, Yam AY-W, Baughman J, Parnot C & Frydman J (2006) Systems analyses reveal two chaperone networks with distinct functions in eukaryotic cells. Cell 124: 75–88

Albuquerque CP, Smolka MB, Payne SH, Bafna V, Eng J & Zhou H (2008) A multidimensional chromatography technology for in-depth phosphoproteome analysis. Mol. Cell Proteomics 7: 1389–1396

Balchin D, Hayer-Hartl M & Hartl FU (2016) In vivo aspects of protein folding and quality control. Science 353: aac4354

Bandhakavi S, Xie H, O’Callaghan B, Sakurai H, Kim D-H & Griffin TJ (2008) Hsf1 activation inhibits rapamycin resistance and TOR signaling in yeast revealed by combined proteomic and genetic analysis. PLoS ONE 3: e1598

Bański P, Kodiha M & Stochaj U (2010) Chaperones and multitasking proteins in the nucleolus: networking together for survival? Trends Biochem. Sci. 35: 361–367

Barbet NC, Schneider U, Helliwell SB, Stansfield I, Tuite MF & Hall MN (1996) TOR controls translation initiation and early G1 progression in yeast. Mol. Biol. Cell 7: 25–42

Becker J, Walter W, Yan W & Craig EA (1996) Functional interaction of cytosolic hsp70 and a DnaJ-related protein, Ydj1p, in protein translocation in vivo. Mol. Cell. Biol. 16: 4378–4386

Beltrao P, Albanèse V, Kenner LR, Swaney DL, Burlingame A, Villén J, Lim WA, Fraser JS, Frydman J & Krogan NJ (2012) Systematic functional prioritization of protein posttranslational modifications. Cell 150: 413–425

Betting J & Seufert W (1996) A yeast Ubc9 mutant protein with temperature-sensitive in vivo function is subject to conditional proteolysis by a ubiquitin- and proteasome-dependent pathway. J. Biol. Chem. 271: 25790–25796

Bishop AC, Ubersax JA, Petsch DT, Matheos DP, Gray NS, Blethrow J, Shimizu E, Tsien JZ, Schultz PG, Rose MD, Wood JL, Morgan DO & Shokat KM (2000) A chemical switch for inhibitor-sensitive alleles of any protein kinase. Nature 407: 395–401

Boorstein WR, Ziegelhoffer T & Craig EA (1994) Molecular evolution of the HSP70 multigene family. J. Mol. Evol. 38: 1–17

Buchberger A, Theyssen H, Schröder H, McCarty JS, Virgallita G, Milkereit P, Reinstein J & Bukau B (1995) Nucleotide-induced Conformational Changes in the ATPase and Substrate Binding Domains of the DnaK Chaperone Provide Evidence for Interdomain Communication. Journal of Biological Chemistry 270: 16903–16910

Bukau B & Horwich AL (1998) The Hsp70 and Hsp60 Chaperone Machines. Cell 92: 351–366

Cashikar AG, Duennwald M & Lindquist SL (2005) A chaperone pathway in protein disaggregation. Hsp26 alters the nature of protein aggregates to facilitate reactivation by Hsp104. J. Biol. Chem. 280: 23869–23875

Cheetham ME & Caplan AJ (1998) Structure, function and evolution of DnaJ: conservation and adaptation of chaperone function. Cell Stress Chaperones 3: 28–36

Chen B, Retzlaff M, Roos T & Frydman J (2011) Cellular strategies of protein quality control. Cold Spring Harb Perspect Biol 3: a004374

Cherkasov V, Hofmann S, Druffel-Augustin S, Mogk A, Tyedmers J, Stoecklin G & Bukau B (2013) Coordination of translational control and protein homeostasis during severe heat stress. Curr. Biol. 23: 2452–2462

Chernoff YO, Lindquist SL, Ono B, Inge-Vechtomov SG & Liebman SW (1995) Role of the chaperone protein Hsp104 in propagation of the yeast prion-like factor [psi+]. Science 268: 880–884

Clerico EM, Tilitsky JM, Meng W & Gierasch LM (2015) How hsp70 molecular machines interact with their substrates to mediate diverse physiological functions. J. Mol. Biol. 427: 1575–1588

Cloutier P & Coulombe B (2013) Regulation of molecular chaperones through post-translational modifications: decrypting the chaperone code. Biochim. Biophys. Acta 1829: 443–454

Conn CS & Qian S-B (2011) mTOR signaling in protein homeostasis: less is more? Cell Cycle 10: 1940–1947

Daugaard M, Rohde M & Jäättelä M (2007) The heat shock protein 70 family: Highly homologous proteins with overlapping and distinct functions. FEBS Lett. 581: 3702–3710

Derkatch IL, Bradley ME, Hong JY & Liebman SW (2001) Prions Affect the Appearance of Other Prions. Cell 106: 171–182

Duennwald ML, Echeverria A & Shorter J (2012) Small Heat Shock Proteins Potentiate Amyloid Dissolution by Protein Disaggregases from Yeast and Humans. PLoS Biology 10: e1001346

Fan C-Y, Lee S & Cyr DM (2003) Mechanisms for regulation of Hsp70 function by Hsp40. Cell Stress Chaperones 8: 309–316

Fan C-Y, Lee S, Ren H-Y & Cyr DM (2004) Exchangeable chaperone modules contribute to specification of type I and type II Hsp40 cellular function. Mol. Biol. Cell 15: 761–773

Flaherty KM, DeLuca-Flaherty C & McKay DB (1990) Three-dimensional structure of the ATPase fragment of a 70K heat-shock cognate protein. Nature 346: 623–628

Flynn G, Chappell T & Rothman J (1989) Peptide binding and release by proteins implicated as catalysts of protein assembly. Science 245: 385–390

Frottin F, Schueder F, Tiwary S, Gupta R, Körner R, Schlichthaerle T, Cox J, Jungmann R, Hartl FU & Hipp MS (2019) The nucleolus functions as a phase-separated protein quality control compartment. Science 365: 342–347

Genest O, Wickner S & Doyle SM (2019) Hsp90 and Hsp70 chaperones: Collaborators in protein remodeling. Journal of Biological Chemistry 294: 2109–2120

Glover JR & Lindquist S (1998) Hsp104, Hsp70, and Hsp40: a novel chaperone system that rescues previously aggregated proteins. Cell 94: 73–82

Greene MK, Maskos K & Landry SJ (1998) Role of the J-domain in the cooperation of Hsp40 with Hsp70. Proc. Natl. Acad. Sci. U.S.A. 95: 6108–6113

Hane F & Leonenko Z (2014) Effect of Metals on Kinetic Pathways of Amyloid-β Aggregation. Biomolecules 4: 101–116

Hartwell LH, Culotti J, Pringle JR & Reid BJ (1974) Genetic control of the cell division cycle in yeast. Science 183: 46–51

Harvey SL, Charlet A, Haas W, Gygi SP & Kellogg DR (2005) Cdk1-Dependent Regulation of the Mitotic Inhibitor Wee1. Cell 122: 407–420

Hasin N, Cusack SA, Ali SS, Fitzpatrick DA & Jones GW (2014) Global transcript and phenotypic analysis of yeast cells expressing Ssa1, Ssa2, Ssa3 or Ssa4 as sole source of cytosolic Hsp70-Ssa chaperone activity. BMC Genomics 15: 194

Holt LJ, Tuch BB, Villén J, Johnson AD, Gygi SP & Morgan DO (2009) Global analysis of Cdk1 substrate phosphorylation sites provides insights into evolution. Science 325: 1682–1686

Huang H, Chen J, Lu H, Zhou M, Chai Z & Hu Y (2017) Two mTOR inhibitors, rapamycin and Torin 1, differentially regulate iron-induced generation of mitochondrial ROS. Biometals 30: 975–980

Hung G-C & Masison DC (2006) N-terminal domain of yeast Hsp104 chaperone is dispensable for thermotolerance and prion propagation but necessary for curing prions by Hsp104 overexpression. Genetics 173: 611–620

Isoda M, Sako K, Suzuki K, Nishino K, Nakajo N, Ohe M, Ezaki T, Kanemori Y, Inoue D, Ueno H & Sagata N (2011) Dynamic Regulation of Emi2 by Emi2-Bound Cdk1/Plk1/CK1 and PP2A-B56 in Meiotic Arrest of Xenopus Eggs. Developmental Cell 21: 506–519

Jacobson T, Priya S, Sharma SK, Andersson S, Jakobsson S, Tanghe R, Ashouri A, Rauch S, Goloubinoff P, Christen P & Tamás MJ (2017) Cadmium Causes Misfolding and Aggregation of Cytosolic Proteins in Yeast. Molecular and Cellular Biology 37: Available at: http://mcb.asm.org/lookup/doi/10.1128/MCB.00490-16 [Accessed September 2, 2019]

Jaiswal H, Conz C, Otto H, Wölfle T, Fitzke E, Mayer MP & Rospert S (2011) The chaperone network connected to human ribosome-associated complex. Mol. Cell. Biol. 31: 1160–1173

Jiang J, Maes EG, Taylor AB, Wang L, Hinck AP, Lafer EM & Sousa R (2007) Structural basis of J cochaperone binding and regulation of Hsp70. Mol. Cell 28: 422–433

Jones GW & Masison DC (2003) Saccharomyces cerevisiae Hsp70 mutations affect [PSI+] prion propagation and cell growth differently and implicate Hsp40 and tetratricopeptide repeat cochaperones in impairment of [PSI+]. Genetics 163: 495–506

Jung G, Jones G, Wegrzyn RD & Masison DC (2000) A role for cytosolic hsp70 in yeast [PSI(+)] prion propagation and [PSI(+)] as a cellular stress. Genetics 156: 559–570

Kaganovich D, Kopito R & Frydman J (2008) Misfolded proteins partition between two distinct quality control compartments. Nature 454: 1088–1095

Kampinga HH & Craig EA (2010) The HSP70 chaperone machinery: J proteins as drivers of functional specificity. Nat. Rev. Mol. Cell Biol. 11: 579–592

Kim S, Schilke B, Craig EA & Horwich AL (1998) Folding in vivo of a newly translated yeast cytosolic enzyme is mediated by the SSA class of cytosolic yeast Hsp70 proteins. Proc. Natl. Acad. Sci. U.S.A. 95: 12860–12865

Kim S-T, Lim D-S, Canman CE & Kastan MB (1999) Substrate Specificities and Identification of Putative Substrates of ATM Kinase Family Members. Journal of Biological Chemistry 274: 37538–37543

Kim YE, Hipp MS, Bracher A, Hayer-Hartl M & Hartl FU (2013) Molecular chaperone functions in protein folding and proteostasis. Annu. Rev. Biochem. 82: 323–355

Kityk R, Kopp J, Sinning I & Mayer MP (2012) Structure and dynamics of the ATP-bound open conformation of Hsp70 chaperones. Mol. Cell 48: 863–874

Koplin A, Preissler S, Ilina Y, Koch M, Scior A, Erhardt M & Deuerling E (2010) A dual function for chaperones SSB-RAC and the NAC nascent polypeptide-associated complex on ribosomes. J. Cell Biol. 189: 57–68

Kose S, Furuta M & Imamoto N (2012) Hikeshi, a nuclear import carrier for Hsp70s, protects cells from heat shock-induced nuclear damage. Cell 149: 578–589

Kravats AN, Hoskins JR, Reidy M, Johnson JL, Doyle SM, Genest O, Masison DC & Wickner S (2018) Functional and physical interaction between yeast Hsp90 and Hsp70. Proc. Natl. Acad. Sci. U.S.A. 115: E2210–E2219

Kusubata M, Matsuoka Y, Tsujimura K, Ito H, Ando S, Kamijo M, Yasuda H, Ohba Y, Okumura E, Kishimoto T & Inagaki M (1993) cdc2 Kinase Phosphorylation of Desmin at Three Serine/Threonine Residues in the Amino-Terminal Head Domain. Biochemical and Biophysical Research Communications 190: 927–934

Lee J, Kim J-H, Biter AB, Sielaff B, Lee S & Tsai FTF (2013) Heat shock protein (Hsp) 70 is an activator of the Hsp104 motor. Proc. Natl. Acad. Sci. U.S.A. 110: 8513–8518

Liebman SW & Chernoff YO (2012) Prions in Yeast. Genetics 191: 1041–1072

Liu Q & Hendrickson WA (2007) Insights into Hsp70 chaperone activity from a crystal structure of the yeast Hsp110 Sse1. Cell 131: 106–120

Matsuoka S, Ballif BA, Smogorzewska A, McDonald ER, Hurov KE, Luo J, Bakalarski CE, Zhao Z, Solimini N, Lerenthal Y, Shiloh Y, Gygi SP & Elledge SJ (2007) ATM and ATR substrate analysis reveals extensive protein networks responsive to DNA damage. Science 316: 1160–1166

Mayer MP (2013) Hsp70 chaperone dynamics and molecular mechanism. Trends Biochem. Sci. 38: 507–514

Mayer MP & Bukau B (1998) Hsp70 chaperone systems: diversity of cellular functions and mechanism of action. Biol. Chem. 379: 261–268

McCusker D, Denison C, Anderson S, Egelhofer TA, Yates JR, Gygi SP & Kellogg DR (2007) Cdk1 coordinates cell-surface growth with the cell cycle. Nature Cell Biology 9: 506–515

Mendenhall MD, Jones CA & Reed SI (1987) Dual regulation of the yeast CDC28-p40 protein kinase complex: cell cycle, pheromone, and nutrient limitation effects. Cell 50: 927–935

Mogk A, Bukau B & Kampinga HH (2018) Cellular Handling of Protein Aggregates by Disaggregation Machines. Molecular Cell 69: 214–226

Narayanaswamy R, Levy M, Tsechansky M, Stovall GM, O’Connell JD, Mirrielees J, Ellington AD & Marcotte EM (2009) Widespread reorganization of metabolic enzymes into reversible assemblies upon nutrient starvation. Proceedings of the National Academy of Sciences 106: 10147–10152

Needham PG & Masison DC (2008) Prion-impairing mutations in Hsp70 chaperone Ssa1: effects on ATPase and chaperone activities. Arch. Biochem. Biophys. 478: 167–174

Oka M, Nakai M, Endo T, Lim CR, Kimata Y & Kohno K (1998) Loss of Hsp70-Hsp40 chaperone activity causes abnormal nuclear distribution and aberrant microtubule formation in M-phase of Saccharomyces cerevisiae. J. Biol. Chem. 273: 29727–29737

O’Neill T, Dwyer AJ, Ziv Y, Chan DW, Lees-Miller SP, Abraham RH, Lai JH, Hill D, Shiloh Y, Cantley LC & Rathbun GA (2000) Utilization of Oriented Peptide Libraries to Identify Substrate Motifs Selected by ATM. Journal of Biological Chemistry 275: 22719–22727

Powers MV, Clarke PA & Workman P (2008) Dual targeting of HSC70 and HSP72 inhibits HSP90 function and induces tumor-specific apoptosis. Cancer Cell 14: 250–262

Rowley A, Johnston GC, Butler B, Werner-Washburne M & Singer RA (1993) Heat shock-mediated cell cycle blockage and G1 cyclin expression in the yeast Saccharomyces cerevisiae. Mol. Cell. Biol. 13: 1034–1041

Rüdiger S, Schneider-Mergener J & Bukau B (2001) Its substrate specificity characterizes the DnaJ co-chaperone as a scanning factor for the DnaK chaperone. EMBO J. 20: 1042–1050

Saibil H (2013) Chaperone machines for protein folding, unfolding and disaggregation. Nat. Rev. Mol. Cell Biol. 14: 630–642

Sanchez Y, Taulien J, Borkovich KA & Lindquist S (1992) Hsp104 is required for tolerance to many forms of stress. EMBO J. 11: 2357–2364

Satterwhite LL (1992) Phosphorylation of myosin-II regulatory light chain by cyclin-p34cdc2: a mechanism for the timing of cytokinesis. The Journal of Cell Biology 118: 595–605

Schirmer EC, Homann OR, Kowal AS & Lindquist S (2004) Dominant Gain-of-Function Mutations in Hsp104p Reveal Crucial Roles for the Middle Region. Molecular Biology of the Cell 15: 2061–2072

Sharma D & Masison DC (2009) Hsp70 structure, function, regulation and influence on yeast prions. Protein Pept. Lett. 16: 571–581

Sharma SK, De los Rios P, Christen P, Lustig A & Goloubinoff P (2010) The kinetic parameters and energy cost of the Hsp70 chaperone as a polypeptide unfoldase. Nat. Chem. Biol. 6: 914–920

Shorter J & Lindquist S (2004) Hsp104 catalyzes formation and elimination of self-replicating Sup35 prion conformers. Science 304: 1793–1797

Shorter J & Lindquist S (2005) Prions as adaptive conduits of memory and inheritance. Nature Reviews Genetics 6: 435–450

Shulga N, Roberts P, Gu Z, Spitz L, Tabb MM, Nomura M & Goldfarb DS (1996) In vivo nuclear transport kinetics in Saccharomyces cerevisiae: a role for heat shock protein 70 during targeting and translocation. J. Cell Biol. 135: 329–339

Sondheimer N & Lindquist S (2000) Rnq1: an epigenetic modifier of protein function in yeast. Mol. Cell 5: 163–172

Sorger PK & Pelham HR (1988) Yeast heat shock factor is an essential DNA-binding protein that exhibits temperature-dependent phosphorylation. Cell 54: 855–864

Soto C (2003) Unfolding the role of protein misfolding in neurodegenerative diseases. Nat. Rev. Neurosci. 4: 49–60

Stone DE & Craig EA (1990) Self-regulation of 70-kilodalton heat shock proteins in Saccharomyces cerevisiae. Mol. Cell. Biol. 10: 1622–1632

Suzuki K, Sako K, Akiyama K, Isoda M, Senoo C, Nakajo N & Sagata N (2015) Identification of non-Ser/Thr-Pro consensus motifs for Cdk1 and their roles in mitotic regulation of C2H2 zinc finger proteins and Ect2. Scientific Reports 5: Available at: http://www.nature.com/articles/srep07929 [Accessed July 27, 2019]

Tamás M, Sharma S, Ibstedt S, Jacobson T & Christen P (2014) Heavy Metals and Metalloids As a Cause for Protein Misfolding and Aggregation. Biomolecules 4: 252–267

Tchounwou PB, Yedjou CG, Patlolla AK & Sutton DJ (2012) Heavy Metal Toxicity and the Environment. In Molecular, Clinical and Environmental Toxicology, Luch A (ed) pp 133–164. Basel: Springer Basel Available at: http://link.springer.com/10.1007/978-3-7643-8340-4_6 [Accessed September 2, 2019]

Trotter EW, Berenfeld L, Krause SA, Petsko GA & Gray JV (2001) Protein misfolding and temperature up-shift cause G1 arrest via a common mechanism dependent on heat shock factor in Saccharomycescerevisiae. Proc. Natl. Acad. Sci. U.S.A. 98: 7313–7318

Truman AW, Kristjansdottir K, Wolfgeher D, Hasin N, Polier S, Zhang H, Perrett S, Prodromou C, Jones GW & Kron SJ (2012) CDK-dependent Hsp70 Phosphorylation controls G1 cyclin abundance and cell-cycle progression. Cell 151: 1308–1318

Verghese J, Abrams J, Wang Y & Morano KA (2012) Biology of the heat shock response and protein chaperones: budding yeast (Saccharomyces cerevisiae) as a model system. Microbiol. Mol. Biol. Rev. 76: 115–158

Walters RW, Muhlrad D, Garcia J & Parker R (2015) Differential effects of Ydj1 and Sis1 on Hsp70-mediated clearance of stress granules in Saccharomyces cerevisiae. RNA 21: 1660–1671

Werner-Washburne M, Stone DE & Craig EA (1987) Complex interactions among members of an essential subfamily of hsp70 genes in Saccharomyces cerevisiae. Mol. Cell. Biol. 7: 2568–2577

Zheng X, Krakowiak J, Patel N, Beyzavi A, Ezike J, Khalil AS & Pincus D (2016) Dynamic control of Hsf1 during heat shock by a chaperone switch and phosphorylation. Elife 5:

Zhu X, Zhao X, Burkholder WF, Gragerov A, Ogata CM, Gottesman ME & Hendrickson WA (1996) Structural analysis of substrate binding by the molecular chaperone DnaK. Science 272: 1606–1614

